# Identification of a convergent spinal neuron population that encodes itch

**DOI:** 10.1101/2023.09.29.560205

**Authors:** Tayler D. Sheahan, Charles A. Warwick, Abby Y. Cui, David A.A. Baranger, Vijay J. Perry, Kelly M. Smith, Allison P. Manalo, Eileen K. Nguyen, H. Richard Koerber, Sarah E. Ross

**Author notes:** Department of Anesthesiology and Perioperative Care, University of California, Los Angeles, Los Angeles, California, USA. Co-first authors (Tayler D. Sheahan, Charles A. Warwick). Correspondence (T.D.S.), (S.E.R.).

## Abstract

Itch is a protective sensation that drives scratching. Although specific cell types have been proposed to underlie itch, the neural circuit basis for itch remains unclear. Here, we used two-photon Ca^2+^ imaging of the dorsal horn to visualize the neuronal populations that are activated by itch-inducing agents. We identify a convergent population of spinal neurons that is defined by the expression of GRPR. Moreover, we discover that itch is conveyed to the brain via GRPR-expressing spinal output neurons that target the lateral parabrachial nucleus. Further, we show that nalfurafine, a clinically effective kappa opioid receptor agonist, relieves itch by inhibiting GRPR spinoparabrachial neurons. Finally, we demonstrate that a subset of GRPR spinal neurons show persistent, cell-intrinsic Ca^2+^ oscillations. These experiments provide the first population-level view of the spinal neurons that respond to pruritic stimuli, pinpoint the output neurons that convey itch to the brain, and identify the cellular target of kappa opioid receptor agonists for the inhibition of itch.

**In brief:** Through population imaging, Sheahan et al. identify a network of neurons in the dorsal horn that is activated by pruritogens and find that kappa opioid receptor signaling inhibits itch through the selective inhibition of GRPR spinoparabrachial neurons.

**Highlights:**

- Itch-inducing agents drive activity in a common population of GRPR-expressing spinal interneurons
- GRPR spinal projection neurons transmit itch from the spinal cord to the brain
- Kappa opioids reduce itch through the inhibition of GRPR spinoparabrachial neurons
- GRPR activation elicits persistent, intrinsic Ca^2+^ oscillations

## Introduction

Itch is an unpleasant sensation that drives organisms to scratch. Itch-inducing stimuli are detected by peripheral sensory neurons that innervate the skin, conveyed to the spinal cord dorsal horn, and finally relayed to the brain where itch can be experienced as a conscious percept. Itch can also be elicited by exogenous application of specific neuropeptides to the spinal cord, including gastrin-releasing peptide (GRP), substance P (SP) and somatostatin^1–9^. These pharmacological findings have implicated several spinal neuron populations in itch, including excitatory spinal neurons that express either the gastrin releasing peptide receptor (GRPR) or neurokinin-1 receptor (NK1R), as well as inhibitory spinal neurons that express the somatostatin receptor (SSTR). However, these receptors are found in different populations of spinal neurons^10,11^, and the manner in which diverse neuropeptides elicit itch remains unclear.

A second outstanding question regarding the spinal circuitry that underlies itch is the identity of the spinal output neurons that convey itch input from the spinal cord to the brain. The existence of an itch-specific population of output neurons has been reported^12^, but remains controversial, since several other studies have found a broad overlap between spinal output neurons that respond to pruritogens and those that respond to algogens^13–18^. Nevertheless, spinal projection neurons that respond to chemical pruritogens such as histamine are thought to reside within the superficial dorsal horn (primarily lamina I), ascend within the anterolateral tract, and target supraspinal structures including the parabrachial nucleus and the thalamus^13,16,19,20^. At a molecular level, pruriceptive spinal output neurons have been shown to express *Tacr1*, the gene encoding NK1R^20,21^. However, most spinal output neurons express NK1R^21–23^, and thus whether a subset of NK1R spinal output neurons comprise a pathway for itch requires further investigation.

Kappa opioid receptor agonists are one of the most effective treatments for chronic itch in humans and these drugs significantly reduce itch behavior in mice and monkeys in response to a variety of distinct pruritogens^2,5,24–27^. Although their site of action remains unclear, the finding that intrathecal delivery of kappa opioid receptor (KOR) agonists is highly effective to reduce scratching in rodents and monkeys suggests that the spinal cord is a likely target^2,5,27^. We therefore reasoned that the application of a KOR agonist would reduce the activity in one or more itch-responsive populations in the spinal dorsal horn which we might be able to identify through population imaging.

Here we used two-photon (2P) Ca^2+^ imaging to identify a convergent population of spinal neurons that responds to itch-inducing peptides and cutaneous pruritogens. As expected based on previous studies, this convergent itch population included GRPR interneurons. Unexpectedly, however, we found that itch was signaled to the brain through GRPR-expressing spinoparabrachial neurons, providing evidence that a specific population of spinal output neurons are involved in relaying itch. Further, we show that KOR agonists selectively target GRPR spinoparabrachial neurons to suppress itch. Finally, we find that GRPR spinal neurons display cell intrinsic Ca^2+^ oscillations. In sum, our findings reveal the population-level organization of the spinal processing of itch as well as a cellular target for the inhibition of itch by KOR agonists.

## Results

### Itch peptides evoke prolonged activity in superficial dorsal horn neurons that parallels scratching behavior

To visualize population-level activity across spinal excitatory neurons in the superficial dorsal horn, we performed 2P Ca^2+^ imaging using an *ex vivo* spinal cord preparation (Figure 1A). We used transgenic mice harboring the *Vglut2-ires-Cre* and *Rosa-lsl-GCaMP6s* (Ai96) alleles to achieve targeted expression of the calcium-sensitive fluorophore GCaMP6s in excitatory neurons (*Vglut2^Cre^;Rosa^GCaMP6s^* mice). The red fluorescent dye, DiI, was injected into the lateral parabrachial nucleus (lPBN), allowing us to identify spinal projection neurons (Figures 1B, 1C). In these experiments, we recorded the activity of excitatory interneurons and DiI labeled spinoparabrachial neurons (SPBNs) across five optical planes (ranging from 0 - 60 µm from the surface of the gray matter) to capture the activity of neurons in lamina I and II (Figure 1D).

**Figure 1.**
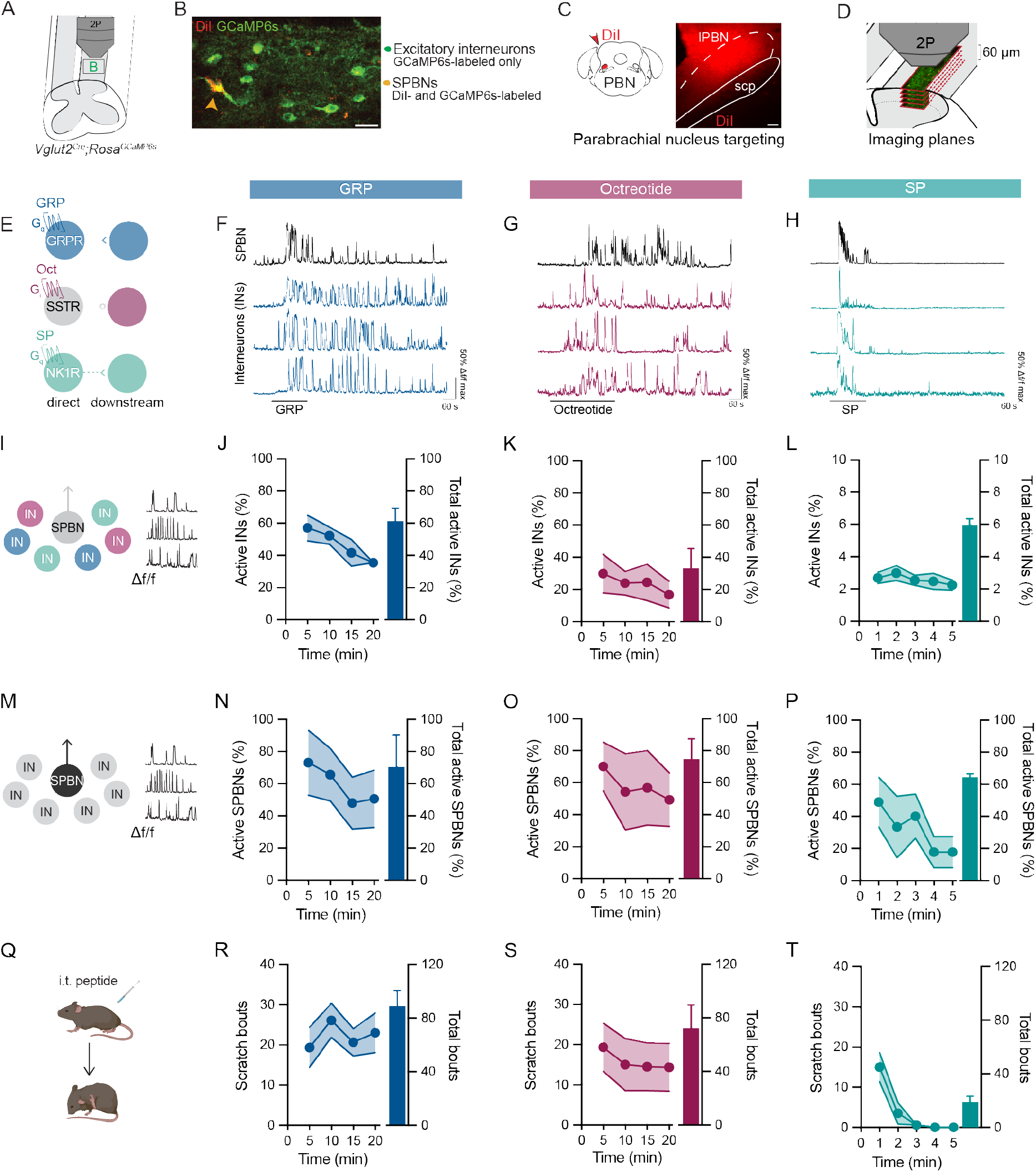
Neuropeptides evoke prolonged activity in spinal neurons that parallels itch behavior. A) 2P Ca^2+^ imaging strategy to visualize the activity of excitatory interneurons and projection neurons within the superficial dorsal horn. B) Representative image of excitatory spinal cord dorsal horn neurons labeled with GCaMP6s (green), including DiI-backlabeled (red) spinoparabrachial neurons (SPBNs, arrowhead). Scale bar, 25 µm. C) Schematic and representative image of targeting the lateral parabrachial nucleus (lPBN) with the retrograde label DiI, Scale bar, 50 µm. D) Schematic of the multiplane Ca^2+^ imaging approach. 5 separate optical planes with a spacing of 15 µm are imaged simultaneously, capturing neurons throughout lamina I and II. E) Schematic of the direct and downstream populations activated in response to bath application of GRP, octreotide, and SP. SSTR is expressed within inhibitory dorsal horn neurons and thus we exclusively recorded neurons that are downstream of SSTR neuron activity. F) Representative ΔF/F Ca^2+^ traces from neurons showing network level responses to 300 nM GRP. G) Representative ΔF/F Ca^2+^ traces from neurons showing network level responses to 200 nM octreotide. H) Representative ΔF/F Ca^2+^ traces from neurons showing network level responses to 1µM SP. I) Schematic showing that the Ca^2+^ activity of excitatory interneurons (INs) was analyzed in panels J-L. J) Percentage of excitatory superficial dorsal horn INs activated by GRP over time (left y-axis) and total percentage of neurons that showed any activity in response to GRP (right y-axis) (n= 213-596 total neurons/mouse, N=3 mice). Data are shown as mean ± SEM. K) Percentage of excitatory superficial dorsal horn INs activated by octreotide over time (left y-axis) and total percentage of neurons that showed any activity in response to octreotide (right y-axis) (n= 213-596 total neurons/mouse, N=3 mice). Data are shown as mean ± SEM. L) Percentage of excitatory superficial dorsal horn INs activated by SP over time (left y-axis) and total percentage of neurons that showed any activity in response to SP (right y-axis) (n= 186-450 total neurons/mouse, N=5 mice). Data are shown as mean ± SEM. M) Schematic showing that the Ca^2+^ activity of SPBNs was analyzed in panels N-P. N) Percentage of SPBNs activated by GRP over time (left y-axis) and total percentage of SPNs that showed any activity in response to GRP (right y-axis) (n= 6-14 SPBNs/mouse, N=3 mice). Data are shown as mean ± SEM. O) Percentage of SPBNs activated by octreotide over time (left y-axis) and total percentage of SPBNs that showed any activity in response to octreotide (right y-axis) (n= 5-14 SPBNs/mouse, N=3 mice). Data are shown as mean ± SEM. P) Percentage of SPBNs activated by SP over time (left y-axis) and total percentage of SPNs that showed any activity in response to SP (right y-axis) (n= 1-14 SPBNs/mouse, N=3 mice). Data are shown as mean ± SEM. Q) Agonists targeting GRPR (295 ng GRP), or SSTR (30 ng octreotide), or NK1R (400 ng SP) were administered intrathecally and spontaneous scratching was quantified in 5-min bins. R) Time course of scratching (left y-axis) and total scratch bouts (right y-axis) following intrathecal injection of GRP (N=15 mice). Data are shown as mean ± SEM. S) Time course of scratching (left y-axis) and total scratch bouts (right y-axis) following intrathecal injection of octreotide (N=8 mice). Data are shown as mean ± SEM. T) Time course of scratching (left y-axis) and total scratch bouts (right y-axis) following intrathecal injection of SP (N=10 mice). Data are shown as mean ± SEM.

GRP, somatostatin (or its analog octreotide), and SP are three neuropeptides that cause scratching when delivered intrathecally^1–9^. We therefore sought to visualize the activity of spinal excitatory neurons that is evoked by the application of these itch-inducing peptides (Figures 1E-1P). We anticipated that GRP would drive activity in excitatory neurons because GRPR is a Gq-coupled receptor that is exclusively expressed in a subset of excitatory neurons in the superficial dorsal horn^4,10,28–31^. In response to bath application of GRP we observed activity in ∼60% of spinal excitatory interneurons in the superficial dorsal horn that lasted for at least 20 minutes (Figures 1F, 1J, S1A). Given its widespread nature, it is likely that this neural activity was due to a combination of GRPR neurons responding directly to GRP as well as neurons that are activated downstream of GRPR neuron activity. Application of octreotide likewise gave rise to prolonged activity, which was observed in ∼30% of excitatory interneurons (Figures 1G, 1K, S1B). Because the somatostatin receptors (SSTRs) are Gi-coupled and they are largely expressed in subsets of inhibitory neurons^11^, the observed activity in excitatory neurons in response to octreotide application is likely indirect, through a mechanism of disinhibition (e.g., downstream activity). Similar to GRP, the application of SP could elicit neural activity in neurons that express NK1R (a Gq-coupled receptor) as well as those activated downstream of NK1R neuron activity. However, in contrast to GRP, SP activated only ∼6% percent of excitatory interneurons, and this activity was brief, lasting ∼ 5 min, (Figures 1H, 1L, and S1C), consistent with previous work showing that NK1R is rapidly internalized following its activation^32,33^. We then evaluated corresponding activity within SPBNs in response to GRP, octreotide, and SP (Figure 1M). For each peptide, the overall time course of activity in spinal output neurons was broadly similar to the time course of activity observed in excitatory interneurons (Figures 1N, 1O, 1P, S1D-S1F).

Because itch elicits scratching, we next examined whether the duration of activity in spinal excitatory interneurons and projection neurons was consistent with the duration of peptide-evoked scratching. Wild-type mice were injected intrathecally with GRP, octreotide, or SP, and scratch bouts were quantified over time (Figure 1Q). As predicted, the activity of superficial dorsal horn neurons observed in 2P Ca^2+^ imaging studies paralleled scratching evoked by each neuropeptide (Figures 1R-1T), consistent with the idea that ongoing spinal neuron activity mediates acute itch behaviors.

### Itch neuropeptides engage a convergent spinal neuron population defined by GRPR expression

The simplest explanation for the observation that spinal application of GRP, octreotide, or SP gives rise to scratching is that all three peptides activate a convergent population of spinal neurons that is involved in mediating itch. Alternatively, each peptide might elicit itch through the activation of independent neuronal populations (Figure 2A). To distinguish between these two models, we applied itch-inducing peptides in series to determine whether they activate common subsets of neurons (Figure 2B). GRP and SP were applied in both the presence or absence of TTX to distinguish between direct responders (e.g., those that express the cognate receptor) and downstream responders (e.g., those whose activity is secondary to the response in GRPR neurons or NK1R neurons) (Figure S2A). The activation of SSTR by octreotide, in contrast, only produces activity in downstream (ds) excitatory neurons via disinhibition, which we refer to as SSTR-ds neurons. In these experiments we recorded from >3000 excitatory neurons, 16% (553/3367 neurons) of which were activated by at least one itch-inducing peptide. A variety of response profiles were observed among the recorded neurons, and neurons were categorized based on these responses. (Figures 2C and S2B). Notably, the order of peptide application did not influence the response profiles that we observed, nor did repeated application of peptides cause tachyphylaxis.

**Figure 2.**
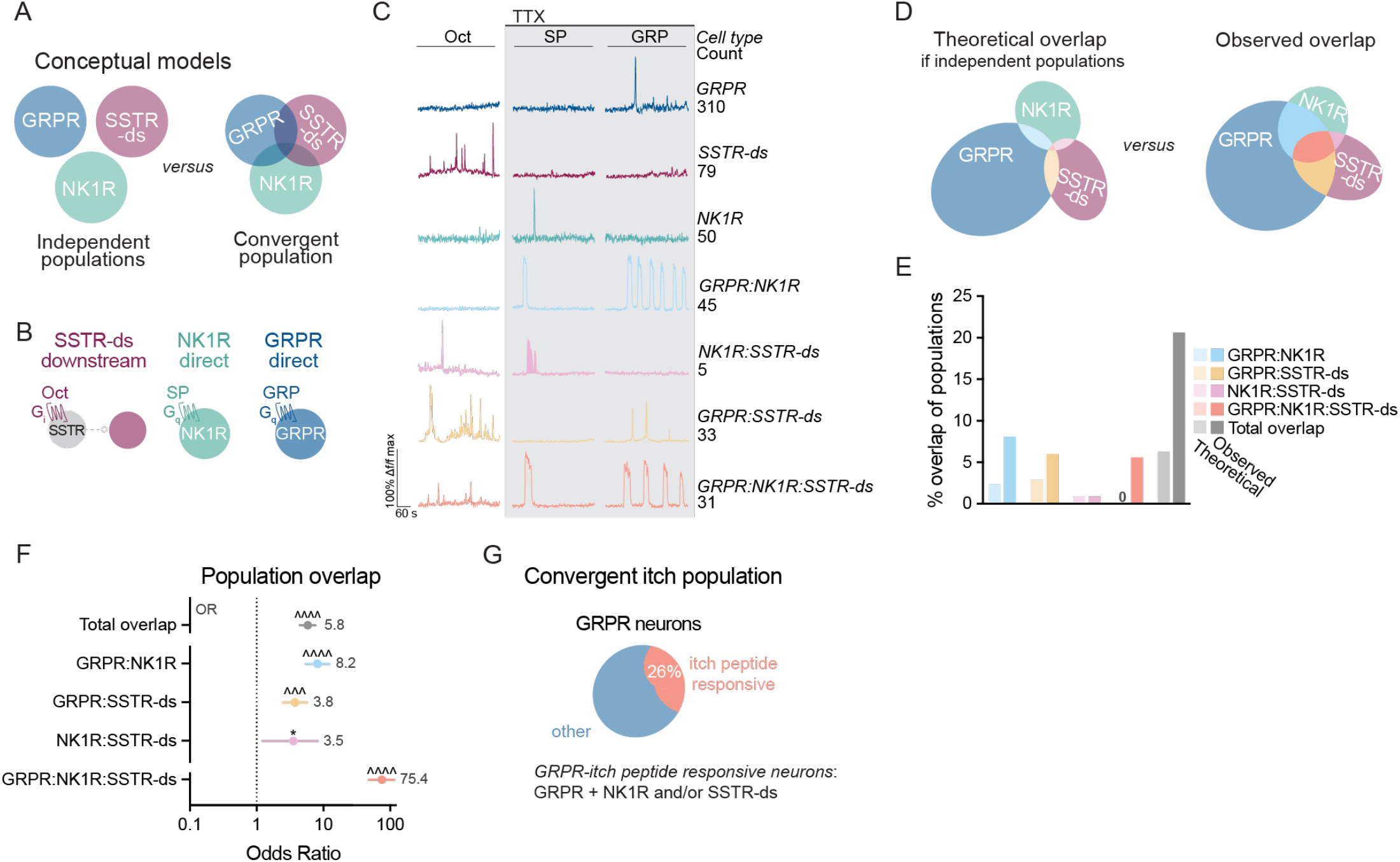
Diverse itch-causing peptides engage a convergent spinal neuron population. A) Models for how diverse itch-causing peptides culminate in scratching behavior. (Left) Different peptides engage independent populations. (Right) Different peptides engage a convergent population of neurons. B) Schematic of cell types visualized upon application of octreotide (200 nM) in the absence of TTX (SSTR-ds) and application of SP (1µM) or GRP (300 nM) in the presence of TTX (500 nM) (NK1R direct, GRPR direct). C) Representative Ca^2+^ imaging traces and cell counts of neurons that responded to one or multiple itch peptides. D) Euler diagram of (left) the extent of population overlap predicted if peptides acted on independent populations of neurons, versus (right) the observed overlap of neurons activated by each peptide. E) Of neurons that responded to itch peptides, the theoretical overlap between populations if populations were independent, compared to the observed overlap between populations. F) GRPR, NK1R, and SSTR-ds populations overlap at frequencies much higher than expected by the prevalence of each cell type (n=3367 total neurons, n=236-599 neurons/mouse, N=8 mice). Odds ratio (OR) analyses, Bonferroni correction for multiple comparisons. OR values for each cell-type in gray. Data are shown as OR estimate ± upper and lower 95% CI. * p < 0.05, ^^^p < 10^-8^, ^^^^p < 10^−10^. G) Schematic showing the convergent itch population of spinal neurons comprised of GRPR neurons that respond to one or more other itch peptides, herein called GRPR itch peptide-responsive neurons. GRPR itch peptide-responsive neurons represent 26% of all GRPR neurons n=109/419 GRPR neurons pooled from N=8 mice).

A preliminary analysis of these response profiles suggested that a large fraction of SSTR-ds neurons were GRPR neurons or neurons that coexpress GRPR and NK1R (GRPR:NK1R neurons). To examine whether this observation could have occurred by chance, we calculated the theoretical overlap if each itch-inducing peptide activated independent populations and compared this with the observed overlap (Figures 2D and 2E). For instance, the theoretical overlap of GRPR, NK1R, and SSTR-ds populations was 0.1%, whereas the observed overlap was 5.6% (Figure 2E). For statistical analysis, we performed a mixed effects log-linear regression to calculate an odds ratio (OR), which reflects whether an observed overlap of populations is larger or smaller than would be expected if the populations were independent. These analyses can be considered an extension of the classic chi-squared test to three or more variables, and further allow us to adjust for differences between mice. We found that the total overlap of cells responding to all three peptides (e.g, GRPR:NK1R:SSTR-ds population) was significantly larger (p = 6.76 x 10^-79^) than predicted if each peptide activated independent populations of neurons (Figures 2E and 2F). Dramatic enrichment was also observed in GRPR:NK1R and GRPR:SSTR-ds populations (Figures 2E and 2F). These results supported the existence of two GRPR spinal neuron populations: one that is broadly responsive to itch-causing peptides, which represent ∼26% of total GRPR neurons, and another that is unresponsive to these peptides (Figure 2G). We similarly observed a common population that was downstream of GRPR, NK1R, and SSTR neurons (Figures S2C and S2D). The finding that common neurons are activated by all three pruritogenic peptides strongly suggests that this convergent population is involved in encoding itch. This convergent population is composed of a subset of GRPR neurons, as well as a common population of neurons that is activated downstream of GRPR itch peptide-responsive neurons.

### GPRR neurons encode responses to cutaneous pruritogens

Our next goal was to visualize the neurons that are activated in response to cutaneous itch, and to determine whether pruritic input from the periphery activates the same convergent population of spinal excitatory neurons that was activated by spinally-applied peptides. For these experiments, we used intradermal compound 48/80 as a model of urticaria (hives)^33^, a condition that is caused by the degranulation of cutaneous mast cells and the release of a variety of endogenous pruritogens including histamine, serotonin, and proteases. To visualize the spinal neurons that respond to intradermal compound 48/80, we used 2P Ca^2+^ imaging of an *ex vivo* somatosensory preparation in which the hairy skin of the dorsal hindpaw through the proximal hip and corresponding nerves (saphenous and lateral femoral cutaneous) are dissected in continuum from the skin through the dorsal root ganglia and to the intact spinal cord (Figures 3A and 3B). As in earlier experiments, excitatory neurons were visualized using *Vglut2^Cre^; Rosa^GCaMP^*^6s^ mice and SPBNs were back labeled with DiI. During imaging sessions, we began by mapping the receptive fields of spinal dorsal horn neurons with punctate mechanical stimuli (Figure 3C). Thereafter, saline (as a control injection) followed by compound 48/80 were injected intradermally. Once the activity evoked by compound 48/80 activity waned, we applied octreotide, GRP, and SP to examine whether these itch-inducing peptides activated the same network of neurons as compound 48/80. Finally, additional pharmacological profiling was performed to identify cell types based on receptor-mediated responses.

**Figure 3.**
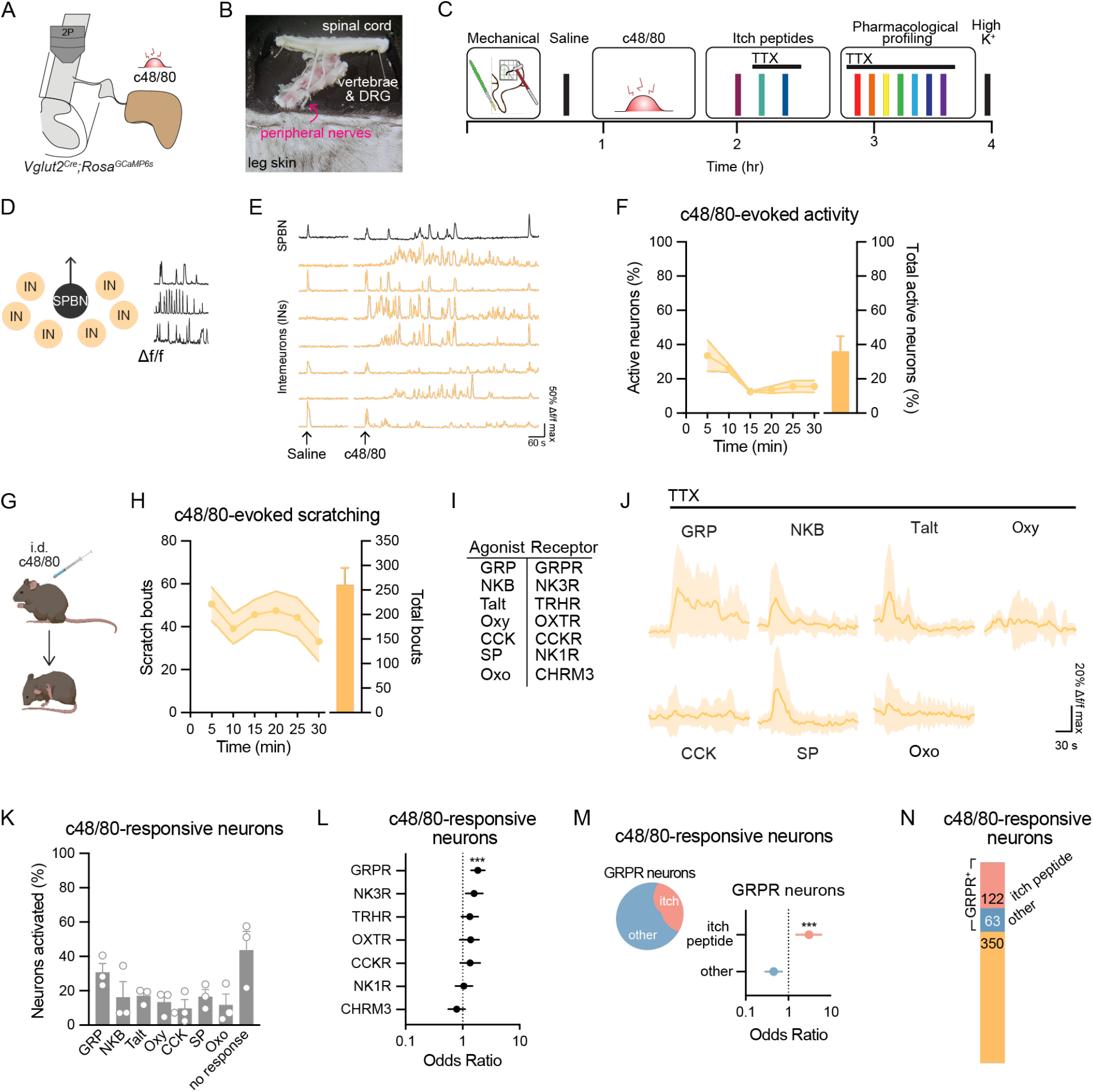
GRPR spinal neurons encode responses to cutaneous pruritogens. A) Experimental strategy for visualizing activity evoked in excitatory superficial dorsal horn neurons caused by intradermal injection of compound 48/80 (91 µg). B) Image of the *ex vivo* semi-intact somatosensory preparation. C) Experimental timeline for visualizing activity evoked by intradermal injection of compound 48/80, itch peptides, as well as pharmacological profiling of excitatory superficial dorsal horn neurons. D) Schematic showing that Ca^2+^ activity of both INs and SPBNs was analyzed in panels E-F. E) Representative ΔF/F Ca^2+^ traces from individual neurons that responded to intradermal injection of compound 48/80. F) Percentage of excitatory superficial dorsal horn neurons activated by compound 48/80 over time (left y-axis), and total percentage of neurons that showed any activity in response to compound 48/80 (right y-axis) (n= 354-610 total neurons/mouse, N=3 mice). Data are shown as mean ± SEM. G) Compound 48/80 (91 µg) was administered intradermally and scratching was quantified in 5-min bins. H) Time course of scratching (left y-axis) and total scratch bouts (right y-axis) following intradermal injection of compound 48/80 (N=11 mice). Data are shown as mean ± SEM. I) Ligands used to pharmacologically profile spinal neuron populations and their cognate Gq-coupled receptors. J) Representative normalized ΔF/F Ca^2+^ traces from experiments pharmacologically profiling compound 48/80-activated neurons. Only neurons that responded to at least 1 ligand were included. Each cell’s ΔF/F trace was normalized to a maximum of 1 and a minimum of 0 (n=44 neurons, N=1 mouse). Data are shown as mean ± SEM. K) Percentage of compound 48/80-responsive neurons that also responded to the application of ligands used for pharmacological profiling (n=85-325 compound 48/80-responsive neurons, N=3 mice). Data are shown as mean ± SEM, with closed circles representing individual mice. The percentage of activated neurons adds up to more than 100% because many neurons responded to more than one ligand. L) GRPR is the only receptor associated with whether a neuron responded to compound 48/80 (n=85-325 compound 48/80-responsive neurons, N=3 mice). Odds ratio (OR) analyses, Bonferroni correction for multiple comparisons. Data are shown as odds ratio estimate ± upper and lower 95% CI. *** p < 0.001. M) GRPR itch neurons are more likely to respond to compound 48/80 than other GRPR neurons. Odds ratio (OR) analysis. Data are shown as odds ratio estimate ± upper and lower 95% CI. *** p < 0.001. N) Quantification of the number of compound 48/80-responsive neurons that are GRPR itch peptide-responsive neurons, other GRPR neurons, or neurons lacking GRPR (n=535 compound 48/80 neurons pooled from N=3 mice).

In response to intradermal saline, we observed a brief response (10-20 s) in ∼80% percent of excitatory neurons. In contrast, compound 48/80 elicited activity in ∼35% of all excitatory superficial dorsal horn neurons, which peaked in 5 minutes and persisted for at least 30 minutes (Figures 3D-3F, S3A), consistent with the duration of compound 48/80-evoked scratching (Figures 3G, 3H). We then used our previously characterized pharmacological cell profiling method^34^ to assign molecular identities to the spinal neuron populations that responded to compound 48/80. The ligands that we used for this profiling allowed us to visualize neurons that express GRPR, NK3R, TRHR, OXTR, CCKR, NK1R, and CHRM3, thereby enabling the identification of 66% of 48/80-responsive neurons (351/535 neurons) in lamina I and IIi (Figures 3I, 3J, 3K, S3B). Among these the ligand that activated the greatest number of compound 48/80 neurons was GRP (Figure 3K), which comprised 31% of the compound 48/80-responsive cells. Because a single neuron can express multiple receptors^34^, we next evaluated which specific receptors were significantly associated with whether a neuron responded to cutaneous injection of compound 48/80. The receptor that was most strongly associated with activity in response to compound 48/80 was GRPR (Figure 3L, S3C), with 57% of all GRPR neurons responding to compound 48/80 (185/325 GRPR neurons). Finally, we analyzed which subset of GRPR spinal neurons responded to compound 48/80. Just as in the itch-inducing peptides data set, the same convergent GRPR itch peptide-responsive spinal neuron population was present (Figure S3D). Notably, GRPR neurons that responded to itch peptides (i.e., octreotide and/or SP) were much more likely to respond to compound 48/80 (Figures 3M and 3N). In sum, these analyses show that peripheral itch input activates the same excitatory convergent spinal neuron population as that activated by spinally-applied peptides, and identify GRPR itch peptide-responsive interneurons as the primary excitatory cell-type that responds to compound 48/80.

### GRPR spinal projection neurons convey chemical itch to the parabrachial nucleus

To examine how itch information is relayed to the brain, we analyzed which population(s) of SPBNs are activated in response to the cutaneous injection of compound 48/80. Compound 48/80 elicited prolonged activity in ∼60% of SPBNs (Figures 4A-4C, S4A), of which 96% could be categorized by their responses to agonists used for pharmacological profiling (Figure 4D). Strikingly, most compound 48/80-responsive SPBNs responded to GRP in the presence of TTX, indicating that GRPR is expressed in SPBNs. Specifically, 78% of compound 48/80-responsive SPBNs were GRPR SPBNS (Figure 4D), and 93% of GRPR SPBNs responded to compound 48/80 (Figure 4E); most of these were also activated by additional itch peptides (Figure 4F). Importantly, the proportion of GRPR SPBNs that responded to compound 48/80 was significantly higher than that predicted by chance. In contrast, although many NK1R neurons were found among compound 48/80-responsive SPBNs, this proportion was not higher than would have been expected by chance (Figure S4B). Together, these data suggest that GRPR SPBNs underlie the transmission of itch input to the brain.

**Figure 4.**
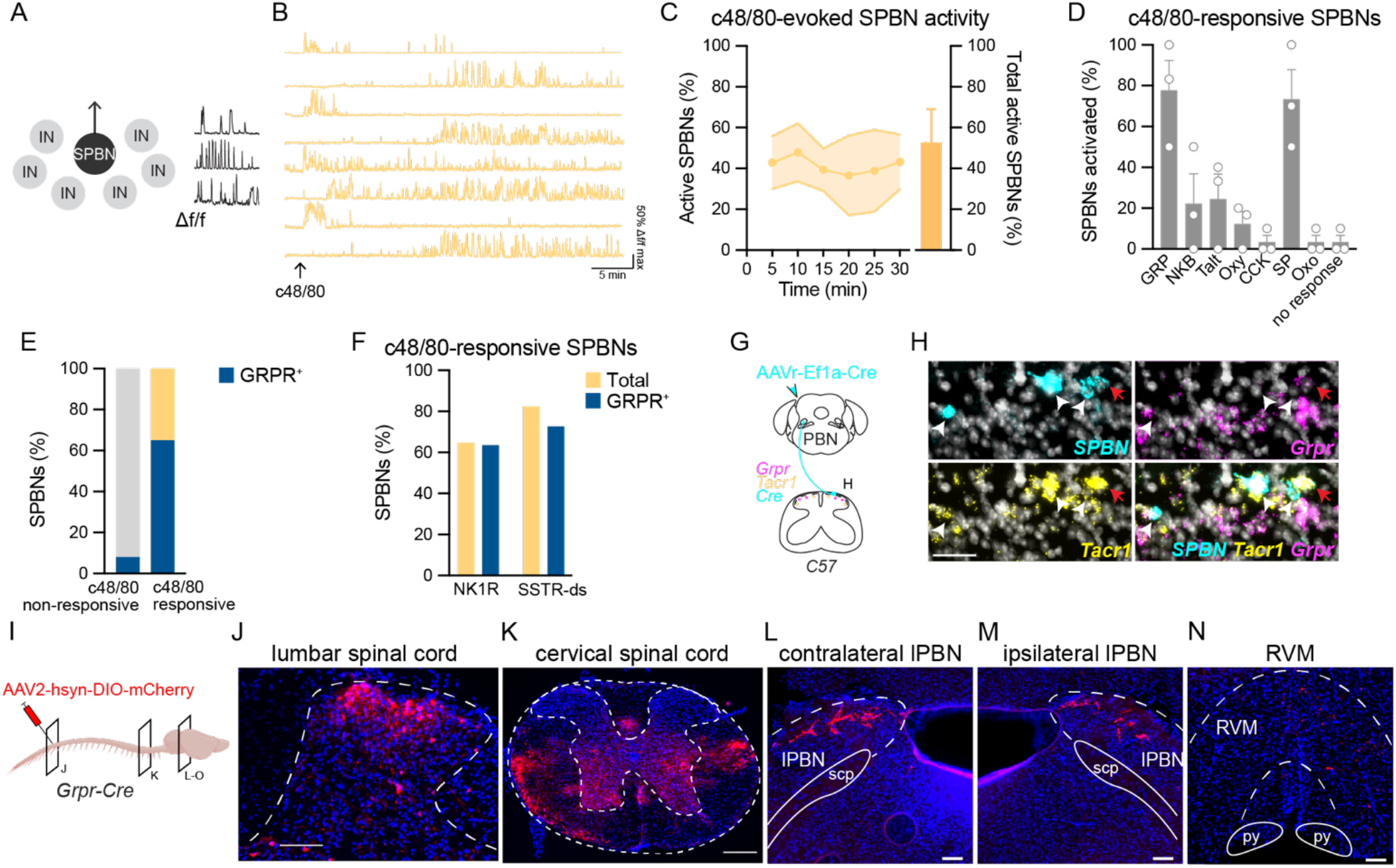
GRPR spinoparabrachial neurons convey itch to the brain. A) Schematic showing that the Ca^2+^ activity of SPBNs was analyzed in panels B-F. B) Representative ΔF/F Ca^2+^ traces from individual SPBNs that responded to intradermal injection of compound 48/80. C) Percentage of SPBNs activated by compound 48/80 over time (left y-axis), and total percentage of SPBNs that showed any activity in response to compound 48/80 (right y-axis) (n= 5-14 PBNs/mouse, N=3 mice). Data are shown as mean ± SEM. D) Percentage of compound 48/80-responsive SPBNs that responded to application of ligands used for pharmacological profiling (n=1-10 compound 48/80-responsive SPBNs, N=3 mice). Data are shown as mean ± SEM, with open circles representing individual mice. The percentage of activated neurons adds up to more than 100% because many neurons responded to more than one ligand. E) Quantification of the number of SPBNs that respond to compound 48/80 and express GRPR and SPBNs that do not respond to compound 48/80 and express GRPR (n=29 SPBNs pooled from N=3 mice). F) GRPR SPBNs that respond to compound 48-80 are GRPR itch peptide-responsive neurons (n=11 GRPR^+^ compound 48/80 responsive SPBNs pooled from N=3 mice). G) Strategy to retrogradely label SPBNs and visualize them with fluorescent *in situ* hybridization (FISH). H) Representative image of FISH of the superficial dorsal horn showing retrogradely labeled SPBNs (Cre, white arrowheads) that express *Grpr*, *Tacr1*, or both transcripts (red arrows). Scale bar, 50 µm. I) Strategy for anterogradely labeling GRPR SPBNs and visualizing their central projections using *Grpr-Cre* mice. (J and K) Representative labeling of *Grpr-Cre* cell bodies at the site of viral injection in the lumbar spinal cord and ascending axons in the cervical spinal cord. Scale bars, 100 µm and 500 µm, respectively. (L-N) Representative images of *Grpr-Cre* spinal projection neuron processes targeting the contralateral lPBN, ipsilateral lPBN, and RVM. PBN: Images representative from 3 of 3 mice analyzed, RVM: Image representative from 2 of 3 mice analyzed. Scale bars, 100 µm.

The observation that GRPR is functionally expressed in SPBNs was surprising as previous evidence suggested GRPR is exclusively expressed in spinal interneurons^29,35^. To explore the possibility that GRPR is expressed in spinal output neurons further, we performed fluorescent *in situ* hybridization on wild-type mice in which SPBNs were retrogradely labeled with virus and probed for expression of *Grpr* (Figure 4G). As a positive control, sections were co-stained with *Tacr1*, the gene encoding NK1R, which was detected in 85% of lamina I SPBNs (Figures 4H, S4C, S4D). *Grpr* was expressed in a subset of lamina I spinal output neurons, and the majority (75%) of *Grpr* SPBNs co-expressed *Tacr1* (Figures 4H, S4C, S4D). Consistent with coexpression of GRPR and NK1R at the mRNA level, we analyzed Ca^2+^ responses of SPBNs to GRP and SP in the presence of TTX. Again, we found strong overlap, with 71% of GRPR SPBNs co-expressing NK1R (Figures S4E, S4F, S4G). Thus, within the dorsal horn, GRPR expression is not restricted to excitatory interneurons; rather GRPR is expressed in two distinct populations: excitatory interneurons, as previously described, and lamina I spinal output neurons.

Lastly, to visualize the central projections of GRPR spinal output neurons, we used *Grpr-Cre* mice and Cre-dependent anterograde labeling through viral expression of AAV2-hsyn-DIO-mCherry in the lumbar spinal cord (Figures 4I and 4J). In these mice, we observed ascending fiber tracts within the anterolateral tract on both the contralateral and ipsilateral sides (Figure 4K). Within the brain, the most prominent targets of *Grpr-Cre* spinal output neurons were the lateral parabrachial nuclei, which were labeled on both the contralateral and ipsilateral sides (Figures 4L and 4M). In a subset of mice, sparse mCherry^+^ projections were also observed in the rostral ventral medulla (RVM) (Figure 4N). This projection pattern suggests that GRPR spinal output neurons are a subset of lamina I spinal output neurons that bilaterally collateralize to the parabrachial nucleus.

### The kappa opioid receptor suppresses itch via inhibition GRPR spinoparabrachial neurons

The kappa agonist nalfurafine is clinically approved in Japan for the treatment of pruritus that accompanies liver or kidney failure^36,37^. Although the specific cell types through which nalfurafine exerts its antipruritic effects remain unknown, the finding that intrathecal nalfurafine reduces numerous types of itch in animal models (e.g., chloroquine-and histamine-evoked itch^5,38^) suggests the involvement of cells in the spinal cord. Consistent with this, we found that mice pretreated with intrathecal nalfurafine scratched significantly less in response to compound 48/80 and chloroquine compared to mice pretreated with vehicle (Figures 5A and 5B). We next asked whether KOR agonism could block scratching in response to application of itch peptides to the spinal cord. Coadministration of nalfurafine significantly attenuated scratching evoked by direct activation of GRPR, SSTR, or NK1R via intrathecal injection of itch peptides (Figures 5C, 5D), demonstrating KOR signaling suppresses behavioral responses to both cutaneous pruritogen and spinal itch peptides.

**Figure 5.**
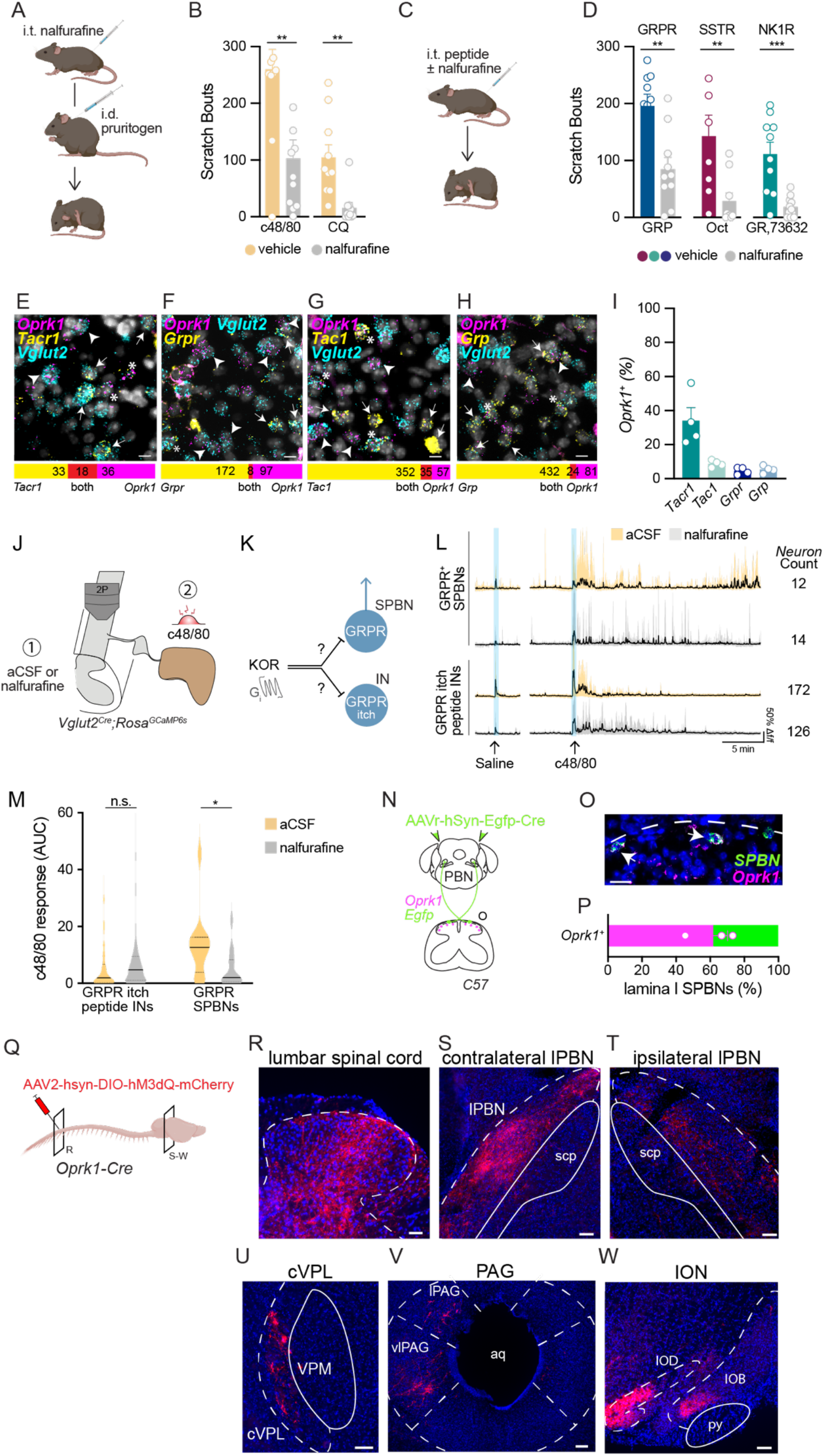
Kappa opioids suppress itch behavior through inhibition of GRPR spinoparabrachial neurons. A) Mice were first pretreated with either intrathecal vehicle or nalfurafine. Twenty minutes later mice received an intradermal injection of chloroquine (100 μg) or compound 48/80 (91 μg) into the nape and the number of site-directed scratch bouts was quantified. B) Intrathecal pretreatment with nalfurafine significantly decreased the number of scratch bouts in response to chloroquine and compound 48/80 compared to vehicle in the 30 min immediately following injection. Student’s T-Test: **p<0.01, n=9-11 mice/group. Data are shown as mean ± SEM, with open circles representing individual mice. C) Agonists targeting GRPR (GRP, 295 ng), SSTR (octreotide, 30 ng), or NK1R (GR,73632, 40 ng) were administered intrathecally alone or in combination with the kappa opioid receptor agonist nalfurafine (40 ng) and spontaneous scratching was quantified. D) Intrathecal nalfurafine significantly reduced scratching evoked by intrathecal GRPR, SSTR, and NK1R agonists in the 30 min immediately following injection. Student’s T-Test: **p<0.01, ***p<0.001, n=8-11 mice/group. Data are shown as mean ± SEM, with open circles representing individual mice. E) Representative image (top) and quantification (bottom) of excitatory superficial dorsal horn neurons (*Vglut2*, cyan) that express *Tacr1* (yellow) and *Oprk1* (magenta). Scale bar, 10 μm. F) Representative image (top) and quantification (bottom) of excitatory superficial dorsal horn neurons (*Vglut2*, cyan) that express *Grpr* (yellow) and *Oprk1* (magenta). Scale bar, 10 μm. G) Representative image (top) and quantification (bottom) of excitatory superficial dorsal horn neurons (*Vglut2*, cyan) that express of *Tac1* (yellow) and *Oprk1* (magenta). Scale bar, 10 μm. H) Representative image (top) and quantification (bottom) of excitatory superficial dorsal horn neurons (*Vglut2*, cyan) that express *Grp* (yellow) and *Oprk1* (magenta). Scale bar, 10 μm. I) Percentage of *Tacr1, Tac1, Grpr,* and *Grp* superficial dorsal horn neurons that coexpress *Oprk1*. Data are shown as mean ± SEM, with open circles representing individual mice. *Tacr1*: n=14-20 neurons/mouse; *Grpr*: n=32-57 neurons/mouse; *Tac1:* n=13-36 neurons/mouse. *Grp* n=18-36 neurons/mouse. N=4 mice for all experiments. J) Experimental approach for examining the effect of nalfurafine on excitatory spinal neuron activity. The spinal cord was first pretreated with either aCSF or nalfurafine (500 nM) for 20 min, followed by an intradermal injection of compound 48/80 (91 µg). Then, as in prior compound 48/80 experiments (Figure 3C), itch peptides were applied, and excitatory superficial dorsal horn neurons were pharmacologically profiled. K) Schematic of two potential sites of action of KOR signaling inhibition of itch: GRPR itch peptide-responsive interneurons (INs), and GRPR spinoparabrachial neurons (SPBNs). L) Representative ΔF/F Ca^2+^ traces showing responses of GRPR SPBNs and GRPR itch peptide-responsive INs to intradermal compound 48/80 following pretreatment with aCSF versus nalfurafine, as well as the number of neurons of subtype per condition (neurons pooled from N=3-4 mice per condition). Blue bars correspond to the brief Ca^2+^ transients evoked by intradermal injection alone. Data shown as mean ± 90th percentile. M) Nalfurafine inhibits the Ca^2+^ responses of GRPR SPBNs to compound 48/80 but does not affect the responses of GRPR itch peptide-responsive INs to compound 48/80 (GRPR^+^ SPBNs: aCSF, n=12 neurons pooled from N=3 mice; nalfurafine, n=14 neurons pooled from N=3 mice. GRPR itch peptide-responsive INs: aCSF, n=172 neurons pooled from N=4 mice; nalfurafine, n=126 neurons pooled from N=4 mice). Linear mixed-effect model, Bonferroni correction for multiple comparisons. * p<0.05. N) Strategy to retrogradely label SPBNs and visualize them with fluorescent in situ hybridization (FISH). O) Representative image of retrogradely labeled SPBNs (green) that coexpress Oprk1 (magenta) in the superficial dorsal horn. Scale bar, 25 μm. P) Percentage of SPBNs in the superficial dorsal horn that coexpress *Oprk1*. Data are shown as mean ± SEM, with open circles representing individual mice. n=19-38 neurons/mouse from N=3 mice. Q) Strategy for anterogradely labeling KOR SPBNs and visualizing their central projections using *Oprk1-Cre* mice. R) Representative labeling of *Oprk1-Cre* cell bodies at the site of viral injection in the lumbar spinal cord Scale bar, 50 µm. S-W) Representative images of *Oprk1-Cre* spinal projection neuron processes targeting the contralateral and ipsilateral lPBN, PAG, contralateral cVPL, and contralateral ION. Images representative from 3 of 3 mice analyzed. Scale bars, 100 µm. IOB, inferior olive, subnucleus B of the medial nucleus; IOD, inferior olive, dorsal nucleus.

Next, to determine whether KOR is expressed within itch spinal neurons, we performed fluorescent *in situ* hybridization. We found *Oprk1* (the gene encoding KOR) expression within both excitatory and inhibitory superficial dorsal horn neurons (Figures S5A and S5B), consistent with recent RNA sequencing datasets^39,40^. In light of these findings, we reasoned that the simplest mechanism by which KOR signaling could inhibit itch is by suppressing activity of excitatory superficial dorsal horn neurons. We found *Oprk1* expression in both *Tacr1* and *Grpr* neurons (Figures 5E, 5F, and 5I). In addition to expression in neurons that express itch neuropeptide receptors, *Oprk1* is also expressed within a subset of neurons that express the itch neuropeptides *Tac1* (substance P) and *Grp* (Figures 5G, 5H, and 5I). Thus, KOR appeared well-positioned to modulate spinal transmission of itch at multiple points within itch spinal circuits, including excitatory interneurons and excitatory spinal projections.

We next sought to pinpoint which spinal neuron populations kappa signaling acts on to suppress itch behavior. Using 2P Ca^2+^ imaging of the *ex vivo* somatosensory preparation, the spinal cord was treated with either aCSF or nalfurafine for 20 minutes prior to intradermal injection of compound 48/80, and activity of excitatory spinal neurons was recorded (Figure 5J). Based on our population-level analyses, two potential sites of action of KOR agonists emerged: GRPR itch peptide-responsive interneurons and GRPR SPBNs (Figure 5K). Surprisingly, nalfurafine reduced the compound 48/80 responses of GRPR SPBNs, but not GRPR itch peptide interneurons (Figures 5L and 5M). Further, nalfurafine did not affect the compound 48/80 responses of SPBNs that lack GRPR, or other GRPR interneurons (Figures S5C-S5E). These findings suggest that kappa opioid signaling selectively suppresses the activity of GRPR SPBNs.

Our finding that KOR agonism suppresses activity of GRPR SPBNs raised the possibility that KOR is expressed in spinal projection neurons. To investigate this, we retrogradely labeled SPBNs with and probed for *Oprk1* (Figure 5N) and found that 60% of lamina I SPBNs express *Oprk1* (Figures 5O and 5P). Finally, we investigated whether *Oprk1* spinal projection neurons target the same brain structures as *Grpr* spinal projection neurons. We visualized the central projections of *Oprk1* spinal output neurons using *Oprk1-Cre* mice and Cre-dependent anterograde labeling through viral expression of AAV2-hsyn-DIO-hM3dQ-mCherry in the lumbar spinal cord (Figures 5Q and 5R). We found *Oprk1-Cre* spinal output neurons had processes within the contralateral and ipsilateral parabrachial nuclei, consistent with the idea that KOR and GRPR are coexpressed in a subset of bilaterally-projecting spinoparabrachial neurons that convey itch to the brain (Figures 5S and 5T). *Oprk1-Cre* labeled spinal output neurons likely include additional populations based on their projections to other brain regions, which included the lateral and ventrolateral periaqueductal gray (PAG), caudal ventral posterior thalamus (cVPL), and the inferior olivary nucleus (Figures 5U-5W). Taken together, these findings suggest that KOR agonists reduce itch through their inhibition of GRPR spinal output neurons.

### A subset GRPR spinal neurons display cell-intrinsic Ca^2+^ oscillations

Throughout our Ca^2+^ imaging studies, we noted that GRP elicited a pattern of Ca^2+^ activity in a subset of spinal neurons that was visually distinct. In these cells, the application of GRP for 3 minutes gave rise to network-level activity that included repeated Ca^2+^ transients that lasted at least 20 minutes, and in one experiment, at least 80 minutes (Figures 6A-6C, S6A-S6C). To distinguish whether these long-lasting oscillations were a consequence of ongoing circuit activity, or whether the oscillations were a cell autonomous phenomenon (Figure 6D), we repeated the GRP application in the presence of TTX. This experiment revealed that GRP-induced oscillations occurred for a similar duration even when network activity was silenced, suggesting that oscillations in GRPR neurons are cell-intrinsic (Figures 6E-6G, S6D-F). To quantify this phenomenon, Ca^2+^ oscillations were defined as 3 or more Ca^2+^ transients with similar amplitudes that occurred at regular intervals in response to brief application of GRP in the presence of TTX (Figure 6H). To determine whether these Ca^2+^ oscillations are specific to GRP, we also tested the effect of other GPCR agonists (Figure 6I). In addition to GRP, both taltirelin (a thyrotropin releasing hormone analog) and SP elicited Ca^2+^ oscillations in excitatory neurons in the dorsal horn; however GRP was more strongly associated with inducing Ca^2+^ oscillations than any other ligand (Figure 6J). Specifically, application of GRP gave rise to persistent Ca^2+^ oscillations in ∼30% of GRP-responsive cells. Because a single neuron can express multiple receptors for these GPCR ligands^34^, we then evaluated whether specific cell-types were associated with Ca^2+^ oscillations. In the case of SP, neurons that express NK1R were only associated with oscillations if they also expressed GRPR (“GRPR:NK1R”, Figure 6K); a similar trend was observed for neurons that oscillated in response to taltirelin and expressed TRHR (“GRPR:TRHR”, Figure 6L). We found that of all of the neurons that displayed cell-intrinsic Ca^2+^ oscillations, GRPR was expressed in 80% of them (197/249 neurons, Figure 6M). Together, these findings suggest that expression of GRPR is a shared feature of the majority of neurons that oscillate, regardless of which agonist causes oscillations.

**Figure 6.**
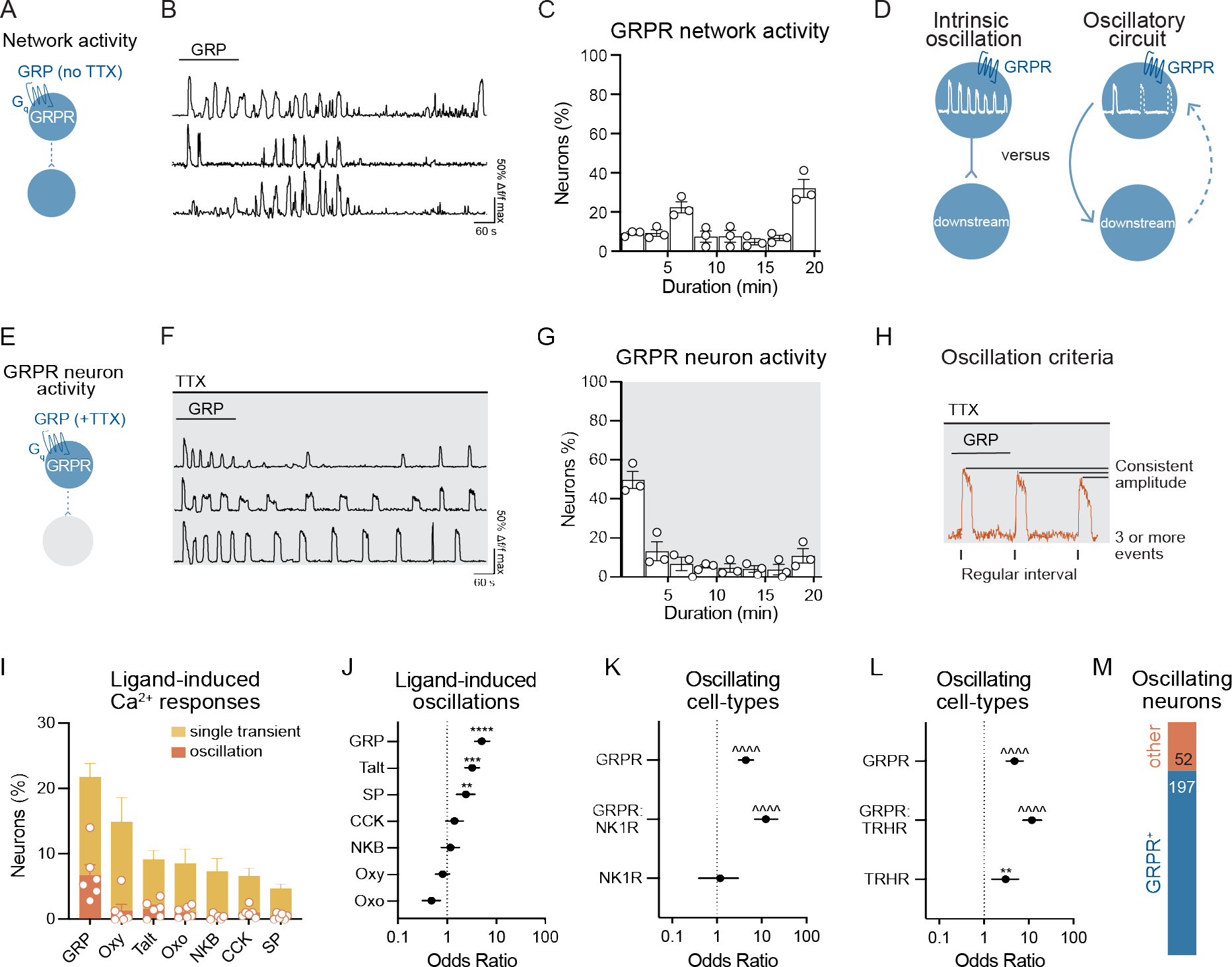
GRPR neurons can display persistent, cell-intrinsic Ca^2+^ oscillations. A) Schematic showing that application of GRP in the absence of TTX produces activity in neurons that express GRPR, as well as neurons that are activated downstream of GRPR neuron activity. B) Representative ΔF/F Ca^2+^ traces from neurons showing GRP-induced repeated Ca^2+^ transients that persist long after GRP is washed out. C) Duration (min) of GRP-evoked network activity (n= 730 total neurons, n=111-447 neurons/mouse, N=3 mice). D) Models for how oscillations could occur in response to GRP. (Left) Oscillations may be an intrinsic feature of GRPR neurons (intrinsic oscillation), or GRPR neurons may form positive feedback loops with downstream neurons to maintain oscillation (oscillatory circuits). E) Schematic showing that application of GRP in the presence of TTX produces activity exclusively in neurons that express GRPR. F) Representative ΔF/F Ca^2+^ traces from neurons showing that GRP application evokes prolonged oscillations in the presence of TTX (500 nM, 8 min pretreatment). G) Duration of GRP-evoked activity in the presence of TTX (n= 229 total neurons, n=65-87 neurons/mouse, N=3 mice). H) Schematic showing criteria for defining Ca^2+^ oscillations. Oscillations were defined as 3 or more Ca^2+^ transients that occur in regular intervals and have consistent amplitudes in the presence of TTX. I) Percentage of neurons that displayed Ca^2+^ transients versus Ca^2+^ oscillations in response to GPCR ligands applied in the presence of TTX (n=275-694 total neurons/mouse from N=6 mice). Data are shown as mean ± SEM, with open circles representing individual mice. The percentage of neurons adds up to more than 100% because many neurons responded to more than one ligand. J) GRP, taltirelin, and SP are all ligands that induce oscillations in neurons (n=101-370 ligand-responsive neurons/mouse, N=6 mice). Odds ratio analysis, Bonferroni correction for multiple comparisons. Data are shown as OR estimate ± upper and lower 95% CI. ** p < 0.01, *** p < 0.001, **** p < 0.000. ^^^^p < 10^−10^. K and L) GRPR expression is the defining feature of cell-types that display cell-intrinsic Ca^2+^ oscillations (n=26-66 oscillating neurons/mouse, N=6 mice). Data are shown as OR estimate ± upper and lower 95% CI. ** p < 0.01, ^^^^ p < 10^−10^. M) GRPR is expressed in the majority of neurons that display cell-intrinsic Ca^2+^ oscillations (n=249 oscillating neurons pooled from N=6 mice).

These findings suggest that a subset of spinal GRPR neurons are notable in their capacity for cell autonomous Ca^2+^ oscillations in response to GRP. The shape and periodicity of these Ca^2+^ oscillations in GRPR neurons were reminiscent of a very well-characterized phenomenon involving cycles of IP3-mediated Ca^2+^ release and SERCA-mediated reuptake (Figure 6SG)^41–43^. Consistent with this mechanism, we found that GRP-induced Ca^2+^ oscillations were abolished in the presence of the IP3 receptor antagonist 2-Aminoethoxydiphenyl borate (2-APB) (Figure 6SH), and triggered spontaneously in the presence of the SERCA pump-inhibitor, CPA (Figure S6I). This type of IP3-mediated Ca^2+^ oscillation has been extensively characterized in a variety of cell types, such as oocytes, pancreatic acinar cells, and osteoclasts^44–46^. Our observations suggest that cell autonomous Ca^2+^ oscillations are also observed in the nervous system, notably in spinal GRPR neurons.

## Discussion

In this study, we visualized the itch-responsive neurons in the dorsal horn at the population-level for the first time. We found that itch-causing agents drive spinal neuron activity that parallels the time course of scratching responses, and that each itch agent we tested recruited a common population of spinal neurons, with expression of GRPR being a defining feature of this itch population. Unexpectedly, we discovered that most of the spinal output neurons that convey itch to the brain are also GRPR neurons, and that these cells are the target of the inhibition of itch by the kappa agonist nalfurafine. Finally, we found that GRPR neurons show a distinctive property in that they are capable of persistent Ca^2+^ oscillations.

### Population coding of itch

Many studies have investigated the contributions of individual cell-types in the dorsal horn to itch, but what has been lacking is a population-level analysis of how itch is encoded. Here, we addressed this fundamental gap in knowledge through 2P Ca^2+^ imaging of the spinal cord dorsal horn. We asked whether three peptides that elicit robust scratching engage a common population of spinal neurons. We found that octreotide acts through disinhibition to drive activity in a population of excitatory neurons that coexpress GRPR and NK1R. These findings are in keeping with and extend the findings of our recent study demonstrating that NK1R is coexpressed within GRPR neurons and that NK1R spinal neurons play a role in itch^4^. Moreover, our analyses revealed that convergence continues with a common population of neurons downstream of GRPR, NK1R, and SSTR neurons. Thus, our study provides novel insight into the mechanisms by which diverse peptides give rise to itch at the population-level. We next visualized the spinal neuron populations activated in response to a cutaneous pruritogen, and found that compound 48/80 recruits the same population of GRPR itch neurons. Together, these data reveal that a common population of neurons underlies scratching that occurs in response to a variety of pruritic agents, both cutaneous and spinal.

In combination with our recent work^34^, these findings also provide novel insights into population coding of pruriceptive versus nociceptive stimuli in the spinal cord. Using the same imaging and pharmacological profiling strategies, we previously delivered the algogen capsaicin intraplantar to identify the cell types that underlie allodynia. In contrast to compound 48/80, capsaicin drove activity in neurons that express NK1R/NK3R (Ex1 neurons), while activity of GRPR neurons (Ex2 neurons) was largely unchanged. Thus, our findings suggest that, although there is some overlap in the spinal neurons that respond to pruritic and algesic stimuli, the primary cell types that is activated by capsaicin and compound 48/80 are distinct.

### GRPR spinal interneurons are heterogeneous

GRPR interneurons neurons have a well-established role in the spinal transmission of itch^1,3,4,21,47–51^. In studies in which GRPR spinal neurons were neurotoxically ablated, itch behaviors were disrupted, while pain behaviors were spared, leading to the idea that GRPR spinal neurons specifically encode itch^47,50^. Our 2P Ca^2+^ imaging studies underscore that GRPR spinal neurons are central to the population coding of itch, as GRPR expression is the defining feature of neurons that belong to the convergent itch population, and GRPR expression is strongly associated whether excitatory neurons were activated by compound 48/80. However, GRPR interneurons neurons were not exclusively found within the convergent itch population, nor did all GRPR interneurons respond to injection of compound 48/80, supporting the idea that GRPR interneurons are a functionally diverse population of neurons. This is in keeping with recent evidence that GRPR is expressed in several different populations of spinal interneurons that may have distinct functions, including itch, noxious heat and mechanical stimuli^30^, as well as mechanical allodynia^52^. Heterogeneity of GRPR neurons is further supported by the detection of *Grpr* in several excitatory dorsal neuron cell-types identified in single cell RNA-seq analyses^10,39^, as well as their spatial distribution throughout the superficial dorsal horn laminae I-IIo^31,34,35,47^. Thus, we propose that there are at least two functional subtypes of GRPR interneurons, of which only one is involved in itch. This framework also extends to NK1R and SSTR neuron populations, which show a broader laminar distribution throughout the superficial dorsal horn^4,53,54^, and have been implicated in central sensitization-induced thermal and mechanical hypersensitivity and gating mechanical pain, respectively^55–58^. In sum, we show that a variety of stimuli that drive scratching do so by recruiting a common population of spinal neurons. Whether this neural organization extends to the manifestation of other aspects of somatosensation is an exciting possibility that merits further investigation.

### GRP elicits persistent Ca^2+^ oscillations

In this study, we report that some neurons in the dorsal horn show recurrent Ca^2+^ oscillations and that the majority of these neurons are GRPR neurons. Ca^2+^ oscillations were originally discovered during fertilization, where the sperm transfers an activating factor, PLCzeta, into oocytes that triggers Ca^2+^ oscillations required for the activation of embryogenesis^1^. Many types of non-excitable cells have been found to oscillate, and these oscillations have been shown to regulate a range of functions including secretion, migration, differentiation, and transcription^2^. For instance, cholecystokinin causes Ca^2+^ oscillations in pancreatic acinar cells, giving rise to the release of digestive enzymes^3^, and ATP causes Ca^2+^ oscillations in osteoclasts, which triggers bone resorption^4^. To our knowledge, our discovery that spinal neurons oscillate is the first report of cell-intrinsic neuronal oscillations in the context of an intact circuit. As found in non-excitable cells, the Ca^2+^ oscillations that we observe in spinal neurons appear to be mediated by repeated cycles of refilling and release of Ca^2+^ from internal stores. Although we do not know the function of these oscillations, we favor the possibility that they may facilitate burst firing in the dorsal horn. Intriguingly, we note that low levels of PLCzeta mRNA are detected in GRPR spinal neurons as well as several other neuronal subtypes across the nervous system^5^, raising the intriguing possibility that PLCzeta-mediated oscillations might be a widespread feature in the nervous system.

### Spinal output neurons for itch

The spinal output neurons that relay itch to the brain have remained largely enigmatic. Chemical itch stimuli have been shown to activate SPBNs that express *Tacr1*^17,20^. However, this classification encompasses the majority of lamina I spinal output neurons^21–23^, and thus whether a distinct subset of these neurons represents the output channel of itch to the brain was unknown. Here, we found the vast majority of SPBNs within the superficial dorsal horn that respond to compound 48/80 express GRPR. Consistent with previous studies^20,21^, these GRPR spinal projection neurons coexpress NK1R, and we further show these are GRPR itch neurons that are activated by other itch peptides like octreotide. The finding that spinal GRPR neurons are composed of both interneurons and spinal output neurons was surprising, as GRPR was believed to be expressed exclusively in interneurons.

In support of this surprising finding, here we provide convergent evidence from fluorescent i*n situ* hybridization studies that GRPR mRNA is expressed in spinal parabrachial neurons. Moreover, by leveraging anterograde labeling strategies, we show that *Grpr-Cre* spinal projection neurons predominantly target the contralateral and ipsilateral lPBN. Consistent with this, a previous study using another genetic allele (*Grpr-Cre^ERT^*^2^) also reported labeling of GRPR spinal neurons in these brain regions^62^. The finding that the spinal projection neurons responsive to chemical itch stimuli appear to specifically target the PBN is particularly interesting when put into the context of the spinal output neurons for mechanical itch, which were recently shown to exclusively target the PBN^20^. Taken together, these findings suggest the PBN serves as the initial point of processing of both chemical and mechanical itch input within supraspinal circuits.

### Kappa opioid receptor inhibition of itch

Here we investigated whether signaling with the spinal cord may underlie KOR inhibition of itch. We and others have previously demonstrated that the endogenous KOR ligand dynorphin is expressed within SSTR spinal neurons^2,5^, but which cells are targeted by dynorphin remained unknown. Because KOR is absent from presumed pruriceptive primary afferents^25,63^, we reasoned that KOR in spinal neurons could be targeted by dynorphin to inhibit scratching. Indeed, our *in situ* hybridization studies suggested *Oprk1* is well-positioned to suppress excitatory spinal neurons that express itch neuropeptides, as well as neuropeptide receptors, including a subset of GRPR neurons. In a previously proposed mechanism for KOR inhibition of itch, KOR was reported to be expressed in roughly half of all GRPR neurons and to inhibit GRPR interneurons through noncanonical PKCδ signaling^38^. However, to our surprise, we found no effect of the KOR agonist nalfurafine on the activity of GRPR interneurons. Rather, we observed that activation of KOR signaling caused robust inhibition of GRPR SPBN activity. These SPBNs may represent the small subset of GRPR spinal neurons inhibited by KOR agonists reported in recent studies^30,64^.

Our imaging data are the first to suggest KOR is expressed in spinal projection neurons. In keeping with our finding that KOR agonism suppresses the activity of GRPR SPBNs, anterograde labeling studies revealed projection patterns suggesting that GRPR and KOR are coexpressed in a population of spinal output neurons that bilaterally target the lateral parabrachial nuclei. Together, these results provide a new mechanism for KOR inhibition of itch in which KOR signaling suppresses the transmission of itch inputs to supraspinal regions. Interestingly, KOR signaling has also been shown to suppress nociceptive behaviors^65^. It is possible that KOR agonists similarly suppress these behaviors by inhibiting the activity of additional spinal output neuron populations, such as those that we found to target the PAG, RVM, and inferior olivary nucleus.

### An updated model for the spinal coding of itch

In summary, this study provides an updated model of the spinal transmission of itch. In this model, a convergent population of interneurons within the superficial dorsal horn that expresses GRPR integrates itch signals that arise from peripheral stimuli, as well as local spinal neuropeptides. Based on evidence that GRPR interneurons arborize in lamina I and make synaptic contacts onto SPBNs^30,35,62^, we propose GRPR interneurons make direct connections with GRPR SPBNs. Our evidence shows inhibition of these SPBNs may underlie spinal KOR inhibition of itch. While we also identified a convergent neuron population downstream of GRPR interneurons that integrates itch inputs, further studies are required to understand where these neurons fit within a model of spinal itch transmission, though one possibility is that they provide inputs onto GRPR spinal output neurons to further shape the coding of sensory inputs to the parabrachial nucleus. More broadly, in this study we shed light on how previously defined spinal cell types are organized at a population level. This approach can be used as a framework in future studies to enable our understanding of the logic of other aspects of spinal somatosensory processing.

## Methods

### Animals

Animals were cared for in compliance with the National Institutes of Health guidelines and experiments were approved by the University of Pittsburgh Institutional Animal Care and Use Committee. *Vglut2-Cre* (Jax, #016963), *RCL-GCaMP6s* (“Ai96”, Jax, #028866), Snap25-2A-GCaMP6s-D (Jax, #025111), and *Grpr-Cre* (Jax, #036668) mouse lines were obtained from Jackson labs. The *Oprk1-Cre* mouse line was previously generated in our lab^69^ (Jax, #035045). All transgenic mouse lines were maintained on a C57BL/6 background. C57BL/6 mice were obtained from Charles River (strain 027). Experiments used a combination of male and female mice. Mice were housed in groups of up to four (males) or five (females) under a 12/12 hr light/dark cycle. Cages were lined with wood chip bedding. Food and water were provided *ab libitum*, and plastic housing domes were provided for enrichment.

### Viruses

The following viruses were used for anterograde and retrograde labeling of SPBNs: AAV2-hsyn-DIO-mCherry (Addgene, #50459-AAV2), AAV2-hsyn-DIO-hM3Dq-mCherry (Addgene, #44361-AAV2), AAVr-Ef1A-Cre (Addgene, #55636-AAVrg), and AAVr-hSyn-eGFP-Cre (Addgene, #105540-AAVrg). Viruses were delivered into the spinal cord or parabrachial nucleus undiluted ranging from titers of 8.6 x 10^10^ to 2.5 x 10^13^.

### Stereotaxic injection and retrograde labeling of spinoparabrachial neurons

Stereotaxic injections were performed on 3.5-5 week old mice for 2P Ca^2+^ imaging experiments and 7-8 week old mice for fluorescent *in situ* hybridization experiments. Mice were anesthetized with 4% isoflurane and maintained in a surgical plane of anesthesia at 2% isoflurane. They were then head-fixed in a stereotaxic frame (Kopf, model 942) and prepared for surgery: the head was shaved, ophthalmic eye ointment was applied, local antiseptic (betadine and ethanol) was applied, and a scalpel was used to make an incision and expose the skull. The skull was leveled using cranial sutures as landmarks. A drill bit (Stoelting 514551) was used to create a burr hole in the skull and a glass capillary filled with a retrograde tracer was lowered through the hole to the injection site. The parabrachial nucleus was targeted at the following coordinates: AP: -5.11 mm, ML ±1.25 mm, DV ±-3.25 mm (from the surface of the skull). A Nanoinject III (Drummond Scientific, 3-000-207) was used to deliver either 250-300 nL of the retrograde tracer Fast DiI oil (2.5 mg/mL) or 500 nL of retrograde viruses at 5 nL/s. The glass capillary was left in place for 5 min following the injection and slowly withdrawn. The scalp was sutured closed with 6-0 vicryl suture. For postoperative care, mice were injected with 5 mg/kg ketofen and 0.03 mg/kg buprenorphine and allowed to recover on a heating pad. Experiments began at least 4 days following DiI injection, and at least 5 weeks after viral injection to allow sufficient time for the retrograde label to reach the spinal cord. To verify targeting of retrograde tracers, brains were collected, post-fixed overnight in 4% paraformaldehyde, sunk in sucrose, and sectioned at 60 µm. Sections were checked for fluorescence in the lateral parabrachial nucleus as mapped in the Allen Mouse Brain Atlas.

### Ex vivo preparations

*Ex vivo* preparations for 2P Ca^2+^ imaging were performed on 5-7 week old mice. All solutions were saturated with 95% O2, 5% CO2 for the duration of the experiment. Sucrose-based artificial cerebrospinal fluid (aCSF) consisted of (in mM): 234 sucrose, 2.5 KCl, 0.5 CaCl2, 10 MgSO4, 1.25 NaH2PO4, 26 NaHCO3, and 11 glucose. Normal aCSF solution consisted of (in mM): 117 NaCl, 3.6 KCl, 2.5 CaCl2, 1.2 MgCl2, 1.2 NaH2PO4, 25 NaHCO3, 11 glucose). Mice were anesthetized with ketamine (87.5 mg/kg)/xylazine (12.5 mg/kg) cocktail prior to beginning dissections. Post dissection, the recording chamber was placed in the multiphoton imaging rig and the sucrose-based aCSF was gradually replaced with normal aCSF over 15 min to avoid shocking the tissue. Meanwhile, the temperature of the aCSF was slowly brought from room temperature to 26**°**C. Imaging began ∼45 min following washout of the recording chamber with aCSF to allow time for recovery of the cells, thermal expansion, and settling of the tissue. Dissection details for each preparation follow below.

### Ex vivo spinal cord preparation

All itch peptide and GRP-induced oscillations experiments were conducted on *Vglut2^Cre^; Rosa^GCaMP^*^6s^ mice. Mice were transcardially perfused with chilled, sucrose-based aCSF. The spinal column was dissected out and chilled in sucrose-based aCSF. The ventral surface of the spinal column was removed, and the spinal cord segment corresponding to the T10 to S1 was carefully removed. Nerve roots were trimmed and the dura and pia mater were removed. Next, using Minutien pins (FST Cat: 26002-202), the spinal cord was secured in a Sylgard-lined recording chamber such that the gray matter of the dorsal horn would be parallel with the imaging objective.

### Ex vivo semi-intact somatosensory preparation

All compound 48/80 control experiments were conducted on *Vglut2^Cre^; Rosa^GCaMP^*^6s^ mice. In kappa agonist experiments, four *Vglut2^Cre^; Rosa^GCaMP^*^6s^ mice and two Snap25*^GCaMP^*^6s^ mice used, with only spinoparabrachial neurons included in analyses from Snap25*^GCaMP^*^6s^ mice. The hindpaw, leg, and back were closely shaved, leaving only 2-3 mm of hair in place. Mice were transcardially perfused with chilled, sucrose-based aCSF. An incision was made along the midline of the back and the spinal cord was exposed via dorsal laminectomy. The spinal column, ribs, and leg were excised and transferred to a Sylgard-lined dish and bathed in sucrose-based aCSF. The lateral femoral cutaneous and saphenous nerves were dissected out in continuum from the skin to the dorsal root ganglia. The thoracolumbar spinal cord was carefully removed from the spinal column, keeping spinal roots intact. Next, the spinal cord was securely pinned with Minutien pins as described above and the dura mater was removed along the entire length of the spinal cord. The pia mater was removed from the L1 - L2 recording area. Finally, the nerves and nerve roots were gently loosened from the tissues, and the skin was pinned out as far from the spinal cord as possible, allowing for a light shield to be placed between the skin and objective at the spinal cord during cutaneous injections.

### Multiphoton imaging

2P Ca^2+^ imaging was performed on a ThorLabs Bergamo II microscope equipped with an 8 kHz resonant-galvo scanpath, GaAsp detectors, a piezo objective scanner for high-speed Z-control during volumetric imaging, and paired with a Leica20X (NA1.0) water immersion lens. A Spectra Physics Mai Tai Ti:Sapphire femtosecond laser tuned to 940 nm was used to excite the fluorophores. Fluorescence was captured with FITC/TRITC emission filter sets with a 570 nm dichroic beam splitter. Five different planes were imaged simultaneously at 8.5 Hz with 15 µm between planes, providing an imaging depth of 0 - 60 µm below the surface of the gray matter. Scanning was performed with a 1.4x optical zoom at a 512 x 256 pixel resolution providing a 514 x 257 µm field of view, yielding a pixel resolution of 1 µm/pixel. This volumetric scanning approach allowed simultaneous sampling from lamina I and II, and yielded between 250-600 excitatory neurons per experiment.

### Receptive field mapping in the ex vivo semi-intact somatosensory preparation

In experiments evaluating cutaneous itch inputs, receptive field mapping was performed under low magnification to empirically identify the region of gray matter receiving primary afferent input from the skin as described previously^34^. Briefly, a wide brush was used as a search stimulus to pinpoint the correct field of view. A wide brush was swept over the entire length of dissected skin, activating as many afferent inputs as possible. The imaging region was refined iteratively until the region with the most fluorescence was in the field of view, and the final 1.4x optical zoom was applied. Next, a 2.0 g von Frey filament was applied to the skin in order to locate the 15 x 15 mm region of skin that received the greatest afferent input, which was marked with a surgical felt tip pen. Imaging, feedback sensors (thermocouples, load cells, etc.), and event times were synchronized using Thosync.

### Intradermal injections in the ex vivo semi-intact somatosensory preparation

Injection of either saline or compound 48/80 (91 µg in 10 µL) was performed with a 31-G needle. Saline was randomized to be given either before or after compound 48/80, and no significant differences were noted in the response. The injection site for both compound 48/80 and saline were centered in the area of skin which elicited the largest response to mechanical stimuli. The needle was held nearly parallel with the skin, bevel side up, and carefully inserted into the skin. An injection was considered successful if it resulted in a visible bleb that did not immediately disappear.

### Drug applications to ex vivo preparations

To measure network-level activity evoked by itch-causing peptides, ligands were bath-applied to the spinal cord. GRP (300 nM) and octreotide (200 nM) were applied for 3 and 7 min, respectively, and subsequent activity was recorded for 20 min. SP (1 µM) was applied to the spinal cord for 3 min, and subsequent activity was recorded for 5 min. To determine whether neurons responded directly to itch-causing peptides and GPCR ligands used for pharmacological profiling, 500 nM TTX (Abcam 120055) was applied to the spinal cord and allowed to circulate for 8-15 min to suppress network activity. Then, agonists were diluted in TTX and applied to the spinal cord for 3 min at the following concentrations: GRP, 300 nM (Tocris 1789); SP, 1 µM (Sigma S6883); NKB, 500 nM (Tocris 1582); taltirelin, 3 µM (Adooq Bioscience A18250); oxytocin, 1µM (Sigma O4375); CCK, 200 nM (Tocris 11661); oxotremorine M, 50 µM (Tocris 1067). The same experimental design was used to determine which GPCR ligands induce cell-intrinsic oscillations. In experiments examining the mechanism of GRP-induced Ca^2+^ oscillations, GRP was applied to the spinal cord to evoke Ca^2+^ oscillations and 2-APB (100 µM, Tocris 1224) 10 min later, followed by a second GRP application after another 10 min. In CPA experiments, the spinal cord was pretreated with CPA (10 µM, Tocris 1235) for 10 min prior to application of GRP and Ca^2+^ responses were recorded for 25 min thereafter. At the end of each experiment, a 30 mM KCl was applied for ∼ 45 sec to confirm cell viability.

### Image processing and data extraction

Recording files were prepared for image processing using macros in FIJI (Image J, NIH). Macros are available on github (https://github.com/cawarwick/ThorStackSplitter). Suite2p (HHMI Janelia) was used for image registration. Then, another ImageJ macro (https://github.com/cawarwick/Suite2p-Output-Processor) was used to review stabilized recordings, generate summary images ROI for selection, and produce averages of the recordings to assist in refining ROIs. We preferred manually selecting ROIs as opposed to using suite2p generated masks because of the heterogeneity of spinal neurons soma size left SPBNs undetected by suite2p. Only cells that had a stable XY location and were free of Z drift were included in analyses. Following ROI selection, FIJI was used to extract raw mean fluorescence intensity. Using either R or Python (Full Python analysis pipeline available on guthub: https://github.com/AbbyCui/CalciumImagingAnalysis), the ΔF/F for each ROI was calculated using the following rolling ball baseline calculation: for each frame (Fi) the 20th percentile value of the surrounding 6.6 min of frames was used as a baseline (Fb) to perform the ΔF/F calculation ((Fi-Fb)/Fb). Rolling ball normalization was applied to minimize effects of gradual photobleaching of GCaMP6s and/or minor changes in ROI location. ΔF/F values for each cell then were plotted over time and traces were visually reviewed. ROIs with recordings that demonstrated Z-drift, were unresponsive to KCl application, or exhibited cell death over the course of the recording were omitted from analysis. Neurons demonstrating basal spontaneous activity were included in counts of total neurons recorded, but were uninterpretable and thus excluded from analyses for responsiveness to itch-inducing neuropeptides or cutaneous stimuli.

### Analysis of Ca^2+^ activity

For analyses of ongoing neuronal activity in response to itch-inducing peptides and compound 48/80, the area under the curve (AUC) of Ca^2+^ activity was calculated for each cell. To determine a response threshold, we first manually identified cells with a visually robust, sustained Ca^2+^ response from a dataset of ∼750 neurons. A receiver operator characteristic analysis was then performed, and the AUC value with 95% specificity was selected to minimize inclusion of false positives. The average AUC threshold was determined by dividing the cumulative AUC threshold by the number of 2.5 min bins (GRP, octreotide, compound 48/80) or 1 min bins (SP) in the recording. This corresponded to an average AUC of at least (0.462 sec*ΔF/F)/bin relative to baseline activity. If cells met this AUC threshold, they were labeled as responders for a given stimulus. For analyses categorizing neurons for convergent population analyses as well as responses to GPCR agonists, neurons were classified as binary responders (yes/no). Neurons were considered binary responders if they displayed a peak Ca^2+^ response within 60 s of applying pharmacological stimuli with an increase in ΔF/F that was both ≥ 30% and ≥ 5 SDs of baseline activity in the 30 s preceding stimulus application. The response window was chosen to account for variability in time required for different drugs to penetrate the spinal cord and their subsequent activation of intracellular signaling cascades.

### Behavior

All behavior experiments were conducted on mice from a C57BL/6J background. Behavior was conducted in a designated, temperature controlled room, with an experimenter blinded to treatment in order to minimize bias. Injection sites were shaved at least 24 hr prior to testing. All behavior experiments were performed during the light cycle between 9 A.M. and 6 P.M. On the day of testing, mice were allotted 30 min to acclimate to a plexiglas chamber on an elevated platform. Behavior platforms had a translucent glass surface, enabling bottom-up recording of animals using a Sony HD Camera (HDR-CX405). The experimenter left the behavior room once the last animal was injected. Behavior was later scored by an observer blind to treatment. Only mice that were successfully injected were included in data analysis.

### Intrathecal neuropeptide-evoked scratching

Itch peptides were administered intrathecally to awake, behaving mice. Mice were pinned by their pelvic girdle and a 25-µL Hamilton Syringe with a 30-G needle attachment was inserted between the L5 and L6 vertebrae. Successful targeting of the intervertebral space was demonstrated by a sudden involuntary lateral tail movement. A total of 5 µL of drug was injected at a rate of 1 µL/s, and the needle was held in place for 5 s prior to removal to minimize backflow. The following peptides were delivered in 5 µL of sterile saline: GRP (295 ng, Tocris 1789), Substance P (400 ng, Sigma S6883), octreotide (30 ng, Tocris 1818), GR,73632 (40 ng, Tocris 1669). Immediately following injection, mice were returned to their plexiglas behavior chambers and recorded for 5-20 min. In studies with kappa opioid receptor agonists, 4% DMSO in sterile saline was used as a vehicle, and itch peptides were either administered alone, or coadministered with nalfurafine (40 ng). Immediately following the injection, mice were returned to their individual plexiglass chamber and were recorded for 30 min.

### Intradermal pruritogen-evoked scratching

Mice were pretreated with either vehicle (4% DMSO), or nalfurafine (40 ng in 5 µL). Twenty minutes later, a 25-µL Hamilton Syringe with a 30-G needle attachment was used to deliver either chloroquine (100 µg in 10 µL, Sigma C6628) or compound 48/80 (91 µg in 10uL, Sigma C2313) intradermally in the nape of the neck. A successful intradermal injection was indicated by the appearance of a bleb at the injection site. Following injection, mice were returned to their Plexiglas behavior chambers and recorded for 30 min.

### RNAscope fluorescent in situ hybridization (FISH)

Animals were anesthetized with isoflurane and quickly decapitated. The L3-L5 spinal cord segments were rapidly removed, flash frozen, and 15 µm sections were directly mounted onto Superfrost Plus slides. FISH experiments were performed according to the manufacturer’s instructions for fresh frozen samples (Advanced Cell Diagnostics, 320293 or 323100). Briefly, spinal cord sections were fixed in ice-cold 4% paraformaldehyde for 15 min, dehydrated in ethanol, permeabilized at room temperature with protease for 15 min, and hybridized at 40**°**C *w*ith gene-specific probes to mouse, including: *Grpr* (#317871), *Tacr1* (#428781), *Cre* (#474001), *Tac1* (#410351), *Grp* (#317861), *Slc17a6* (#456751), *Egfp* (#400281), and *Oprk1* (#316111). Hybridized probe signal was then amplified and fluorescently labeled. Slides were mounted with Prolong Gold with DAPI to visualize nuclei.

### Intraspinal injections

Intraspinal viral injections were performed on 5-7 week old mice. Animals were anesthetized with a ketamine (95 mg/kg)/xylazine(4.8 mg/kg)/acepromazine cocktail (0.95 mg/kg). They were then prepared for surgery: the back was shaved, local antiseptic (betadine and ethanol) was applied to the skin, and a scalpel was used to make an incision over the T12-L3 vertebrae. Muscle and fascia were removed, exposing the L3/L4 and L4/L5 spinal segments. For each segment, a glass capillary filled with virus was lowered 300 µm below the surface of the spinal cord to target the dorsal horn, and a Nanoinject III was used to deliver 500 nL of virus at 5 nL/s. The glass capillary was left in place for 5 min following each injection and slowly withdrawn. The skin was sutured closed with 6-0 vicryl suture. For postoperative care, mice were injected with 5 mg/kg ketofen and 0.03 mg/kg buprenorphine and allowed to recover on a heating pad. Histology experiments began at least 4 weeks following viral injection to allow sufficient time for the viral labeling to reach the brain.

### Immunohistochemistry

For histology studies visualizing the central projection of spinal output neurons, mice were anesthetized with intraperitoneal injection urethane (4 mg/kg, i.p.) and transcardially perfused with 4% paraformaldehyde. Spinal cord and brain were removed and post-fixed in 4% paraformaldehyde for either 2 hr (spinal cord) or overnight (brain). Tissues were then washed in PBS with 0.3% triton (PBS-T), sunk in 30% sucrose, and cryosectioned at either 20 µm (spinal cord) or 40 µm (brain). Sections were incubated in a blocking solution made of 10% donkey serum (Jackson ImmunoResearch, 017-000-121) and 0.3% triton in PBS for 1 hr at room temperature. Sections were incubated overnight at 4**°**C with the primary antibody rabbit anti-RFP (Rockland, 600-401-379) diluted at 1:1K in an antibody buffer consisting of 5% donkey serum and 0.3% triton in PBS. Following washes in PBS-T, sections were incubated for 1 hr at room temperature with the secondary antibody donkey anti-rabbit Alexa Fluor 555 (Invitrogen, A-31572) at 1:500 diluted in antibody buffer. Tissues were washed in PBS and mounted with Prolong Gold with DAPI (Invitrogen, P36931).

### Image acquisition and quantification

Full-thickness tissue sections were imaged using an upright epifluorescent microscope (Olympus BX53 with UPlanSApo 4x, 10x or 20x objective) or confocal microscope (Nikon A1R with an 20x oil-immersion objective). Image analysis was performed off-line using FIJI imaging software (Image J, NIH). In FISH experiments evaluating markers in superficial dorsal horn neurons, the superficial dorsal horn was defined as the region between the surface of the gray matter and the bottom of the substantia gelatinosa, a region ∼ 65 µm thick corresponding to lamina I and II. For FISH experiments evaluating coexpression of *Oprk1* in spinal cord neuronal markers, 3 spinal cord hemisections were manually quantified from 3-4 mice per marker. For FISH experiments visualizing retrogradely labeled SPBNs, 9-15 hemisections per mouse were imaged to ensure enough sparsely labeled projection neurons were available for analysis. In immunohistochemistry experiments assessing the central targets of *Grpr-Cre* and *Oprk1-Cre* spinal projection neurons, brain sections from 3 mice were imaged and brain structures with mCherry/RFP signal were recorded.

### Quantification and statistical analysis

Microsoft Excel, GraphPad Prism, EulerAPE^66^, R, and Python packages including Scipy, numpy, pandas, matplotlib, and other custom code were used for data organization, processing, visualization, and statistical analyses. Log-linear analyses were used to test whether populations of neurons overlap more than expected by chance. Differences between mice were adjusted for using a mixed-effect model (i.e., a random intercept for mouse) when data were available for at least six mice. Otherwise, models included fixed-effects to adjust for the effect of each mouse. Analyses of neuronal responses to GPCR agonists were run as logistic (i.e., compound 48/80 binary responder) and linear (AUC) regressions, adjusting for mouse as fixed-effects. Linear mixed-effect model analyses were used to examine the effect of nalfurafine on Ca^2+^ responses. To reduce the influence of outliers, AUC values were winsorized to 3 SDs and log transformed, and differences between mice were adjusted for using a random intercept for mouse. Throughout the figures, statistical significance is indicated with the following symbols: *p < 0.05, **p < 0.01, ***p < 0.001, ****p < 0.0001, ^^^p < 1 x 10^-8^, ^^^^p <1 x 10^-8^. A Bonferroni correction was used to correct for multiple comparisons when appropriate. Experiment-specific details can be found in the figure legends. Data are presented as mean ± SEM, with a few exceptions. Odds ratio analyses are presented as the odds ratio estimate ± upper and lower 95% CI. Linear regression analyses are presented as estimate ± upper and lower 95% CI. Representative traces of the Ca^2+^ responses of populations of neurons are presented as either median ± interquartile range or mean ± 90th percentile.

| Key resources table |  |  |
| --- | --- | --- |
| REAGENT or RESOURCE | SOURCE | IDENTIFIER |
| Antibodies |  |  |
| rabbit α-RFP | Rockland | Cat#600-401-379; RRID: AB_2209751 |
| donkey α-rabbit Alexa Fluor 555 | Invitrogen | Cat#A-31572; RRID: AB_162543 |
| Bacterial and virus strains |  |  |
| AAV2-hSyn-DIO-mCherry | Addgene | 50459-AAV2 |
| AAV2-hSyn-DIO-hM3Dq-mCherry | Addgene | 44361-AAV2 |
| AAVr-Ef1A-Cre | Addgene | 55636-AAVrg |
| AAVr.hSyn.HI.eGFP-Cre.WPRE.SV40 | Addgene | 105540-AAVrg |
| Chemicals, peptides, and recombinant proteins |  |  |
| Compound 48/80 | Sigma-Aldrich | Cat#C2313; CAS: 848035-21-2 |
| Chloroquine Diphosphate Salt | Sigma-Aldrich | Cat#C6628; CAS: 50-63-5 |
| FAST Dil<sup>TM</sup> oil | Invitrogen | Cat#D3899 |
| Octreotide | Tocris Bioscience | Cat#1818; CAS: 83150-76-9 |
| GRP | Tocris Bioscience | Cat#1789; CAS: 93755-85-2 |
| Substance P | Sigma-Aldrich | S6883; CAS: 137348-11-9 |
| Taltirelin (Talt) | Adooq Bioscience | Cat#A18250; CAS: 103300-74-9 |
| Oxytocin (Oxy) | Sigma-Aldrich | O4375; CAS: 50-56-6 |
| Oxotremorine (Oxo) | Tocris Bioscience | Cat#1067; CAS: 3854-04-4 |
| Cholecystokinin Octapeptide, sulfated (CCK) | Tocris Bioscience | Cat#11661 |
| Neurokinin B (NKB) | Tocris Bioscience | Cat#1908; CAS: 87096-84-2 |
| Tetrodotoxin citrate (TTX) | Abcam | Cat#ab120055; CAS: 18660-81-6 |
| GR, 73632 | Tocris Bioscience | Cat#1669; CAS: 133156-06-6 |
| Nalfurafine Hydrochloride | Adooq Bioscience | Cat#A12579; CAS: 152658-17-8 |
| 2-Aminoethoxydiphenyl borate (2-APB) | Tocris Bioscience | Cat#1224; CAS: 524-95-8 |
| Cyclopiazonic acid (CPA) | Tocris Bioscience | Cat#1235; CAS: 18172-33-3 |
| Critical commercial assays |  |  |
| RNAscope<sup>TM</sup> Probe- Mm-Grpr | ACDbio | Cat#317871 |
| RNAscope<sup>TM</sup> Probe- Mm-Tacr1 | ACDbio | Cat#428781 |
| RNAscope<sup>TM</sup> Probe- Cre | ACDbio | Cat#474001 |
| RNAscope<sup>TM</sup> Probe- Mm-Slc17a6 | ACDbio | Cat#456751 |
| RNAscope<sup>TM</sup> Probe- Mm-Tac1 | ACDbio | Cat#410351 |
| RNAscope<sup>TM</sup> Probe- Mm-Grp | ACDbio | Cat#317861 |
| RNAscope<sup>TM</sup> Probe- Mm-Oprk1 | ACDbio | Cat#316111 |
| RNAscope<sup>TM</sup> Probe- EGFP | ACDbio | Cat#400281 |
| Experimental models: Organisms/strains |  |  |
| Mouse: C57BL6/J | Charles River | Cat: 027 |
| Mouse: *Vglut2-Cre* | Vong et al.<sup>67</sup> | Jax Stock #016963 |
| Mouse: *Ai96 (RCL-GCaMP6s)* | Madisen et al.<sup>68</sup> | Jax Stock #028866 |
| Mouse: *Snap25-2A-GCaMP6s-D* | Madisen et al.<sup>68</sup> | Jax Stock #025111 |
| Mouse: *Grpr-Cre* | Munanairi et al.<sup>38</sup> | Jax Stock #036668 |
| Mouse: *Oprk1-Cre* | Cai et al.<sup>69</sup> | Jax Stock #035045 |
| Software and algorithms |  |  |
| PRISM 9.0 | GraphPad | https://www.graphpad.com/scientific-software/prism/ |
| FIJI | ImageJ | https://www.nature.com/articles/nmeth.2019 |
| suite2p | v0.10.0, HHMI Janelia | https://www.suite2p.org/ |
| R | R Core Team (2021)<sup>70</sup> | https://www.r-project.org/ |
| Python | Python 3.10.7 | https://www.python.org/ |
| EulerAPE | Micallef and Rodgers<sup>66</sup> | https://www.eulardiagrams.org/eulerAPE/ |

## Acknowledgements

We thank all members of the Ross lab for their comments and suggestions, as well as Michael C. Chiang and Haichao C. Chen for their technical assistance. Select figure panels were created using icons from BioRender.

## Inclusion and Diversity

We support inclusive, diverse, and equitable conduct of research.

