## Supplemental Figures for "Identification of a convergent spinal neuron population that encodes itch"

### Supplemental Data

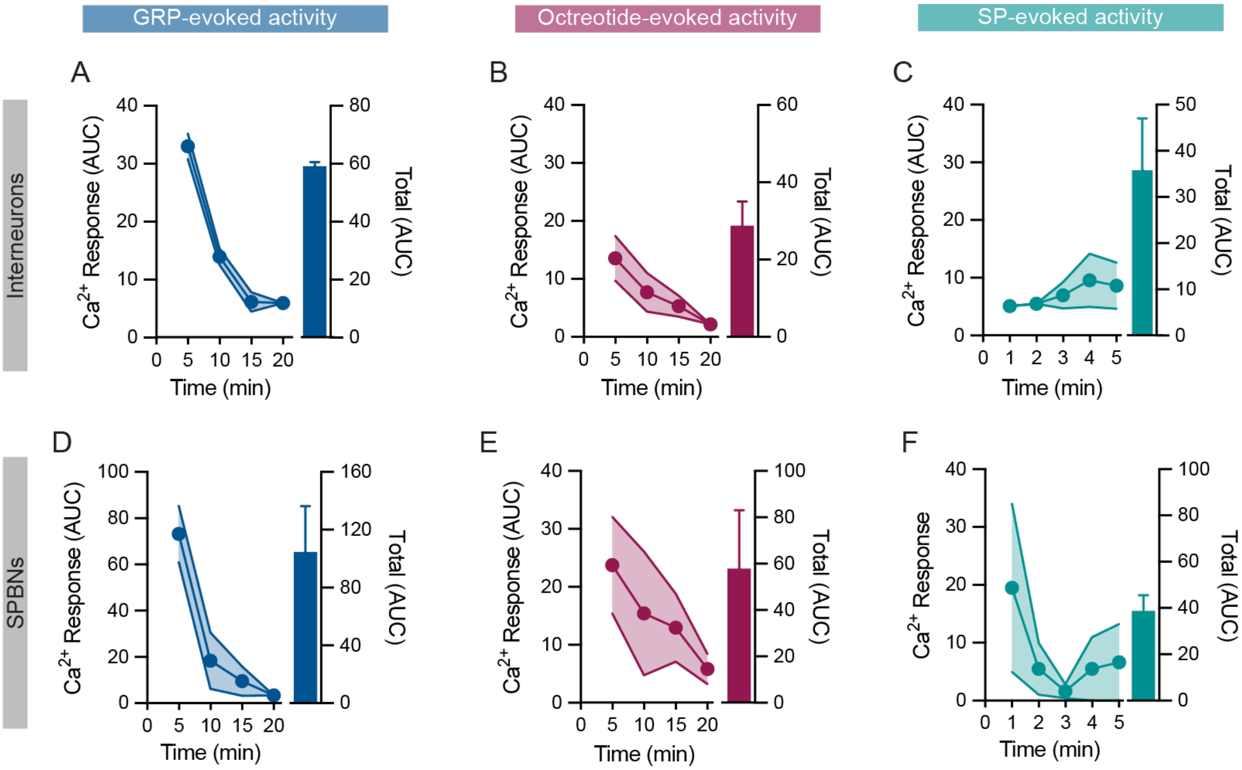


**Supplemental Figure 1**

S1A) AUC over time (left y-axis) and cumulative AUC (right y-axis) in excitatory superficial dorsal horn INs following application of GRP (n=120-460 GRP-responsive neurons/mouse, N=3 mice). Data are shown as mean ± SEM.

S1B) AUC over time (left y-axis) and cumulative AUC (right y-axis) in excitatory superficial dorsal horn INs following application of octreotide (n=59-193 octreotide -responsive neurons/mouse, N=3 mice). Data are shown as mean ± SEM.

S1C) AUC over time (left y-axis) and cumulative AUC (right y-axis) in excitatory superficial dorsal horn INs following application of SP (n=12-44 SP-responsive neurons/mouse, N=5 mice). Data are shown as mean ± SEM.

S1D) AUC over time (left y-axis) and cumulative AUC (right y-axis) in SPBNs following application of GRP (n=2-12 GRP-responsive SPBNs/mouse, N=3 mice). Data are shown as mean ± SEM.

S1E) AUC over time (left y-axis) and cumulative AUC (right y-axis) in SPBNs following application of octreotide (n=3-9 octreotide-responsive SPBNs/mouse, N=3 mice). Data are shown as mean ± SEM.

S1F) AUC over time (left y-axis) and cumulative AUC (right y-axis) in SPBNs following application of SP (n=2-3 SP-responsive SPBNs/mouse, N=3 mice). Data are shown as mean ± SEM.

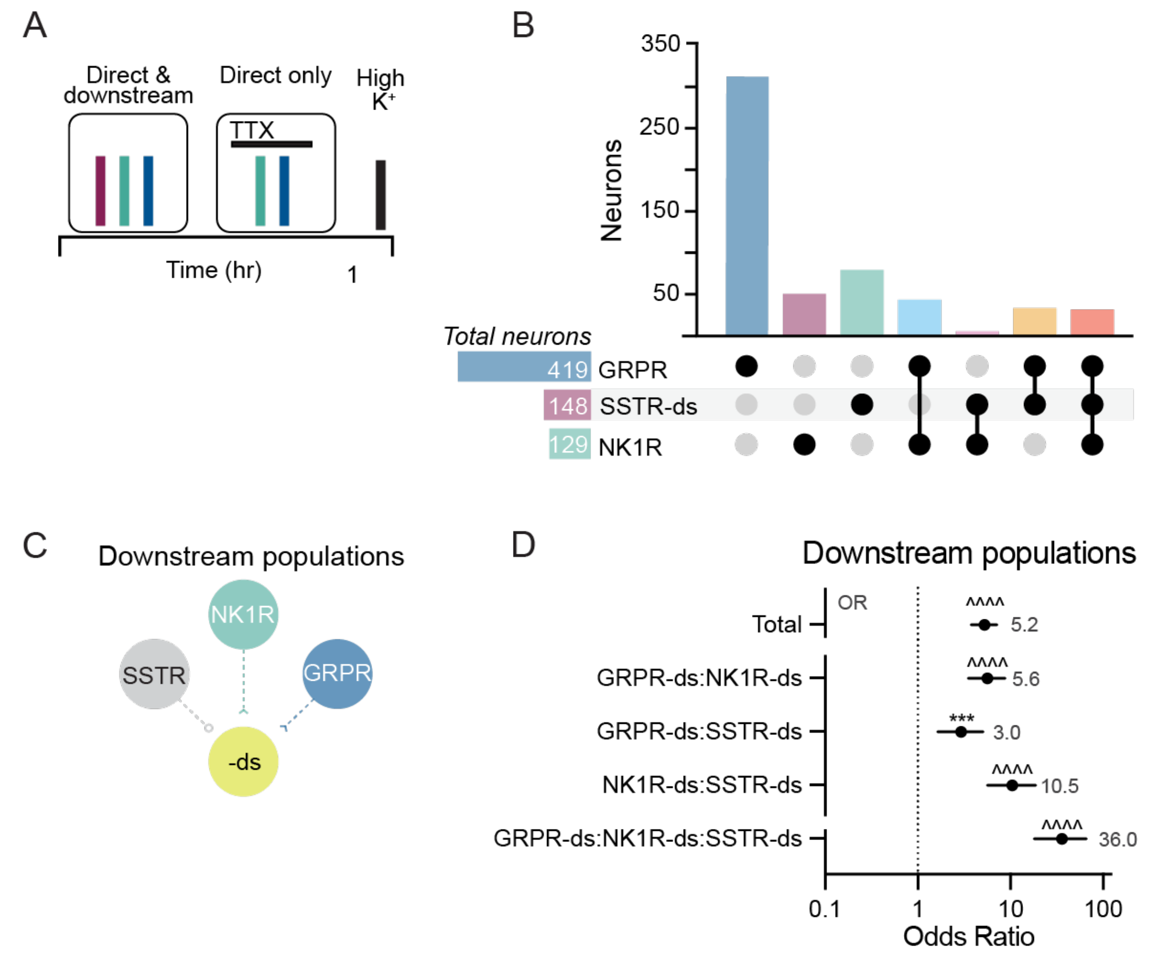


**Supplemental Figure 2**

S2A) Experimental timeline for visualizing activity evoked by the application of itch-inducing peptides in the absence and presence of TTX. A direct responder was defined as a neuron that responded to GRP or SP in both the presence and absence of TTX. In contrast, a downstream responder (e.g., a neuron whose activity is secondary to the response in GRPR neurons or NK1R neurons), was defined as neurons that responded to GRP, SP, or octreotide exclusively in the absence of TTX. A final application of high $K^+$ (30 mM KCl) was applied to visualize all excitatory superficial dorsal horn neurons.

S2B) Number of neurons that responded to one or multiple itch-inducing peptides.

S2C) Schematic showing a common neuron downstream of SSTR, NK1R, and GRPR neurons. In these experiments, we noted that the earliest point of convergence of GRP, SP, and octreotide signaling is on neurons that express both GRPR and NK1R and are downstream of inhibitory SSTR neurons. In light of this, we reasoned that there might be an additional point of convergence: a common population that is downstream of the GRPR:NK1R:SSTR-ds population. To investigate this idea, we performed analogous statistical analyses to ask whether the overlap of GRPR-ds, SSTR-ds, and NK1R-ds neuron populations was greater than that expected by chance.

S2D) GRPR-ds, NK1R-ds, and SSTR-ds populations overlap at frequencies much higher than expected by the prevalence of each cell type. (n=3367 total neurons, n=236-599 neurons/mouse, N=8 mice) Odds ratio (OR) analyses, Bonferroni correction for multiple comparisons. OR values for each cell type are in gray. Data are shown as OR estimate ± upper and lower 95% CI. *** $p < 0.001$, ^^^^$p < 10^{-10}$.

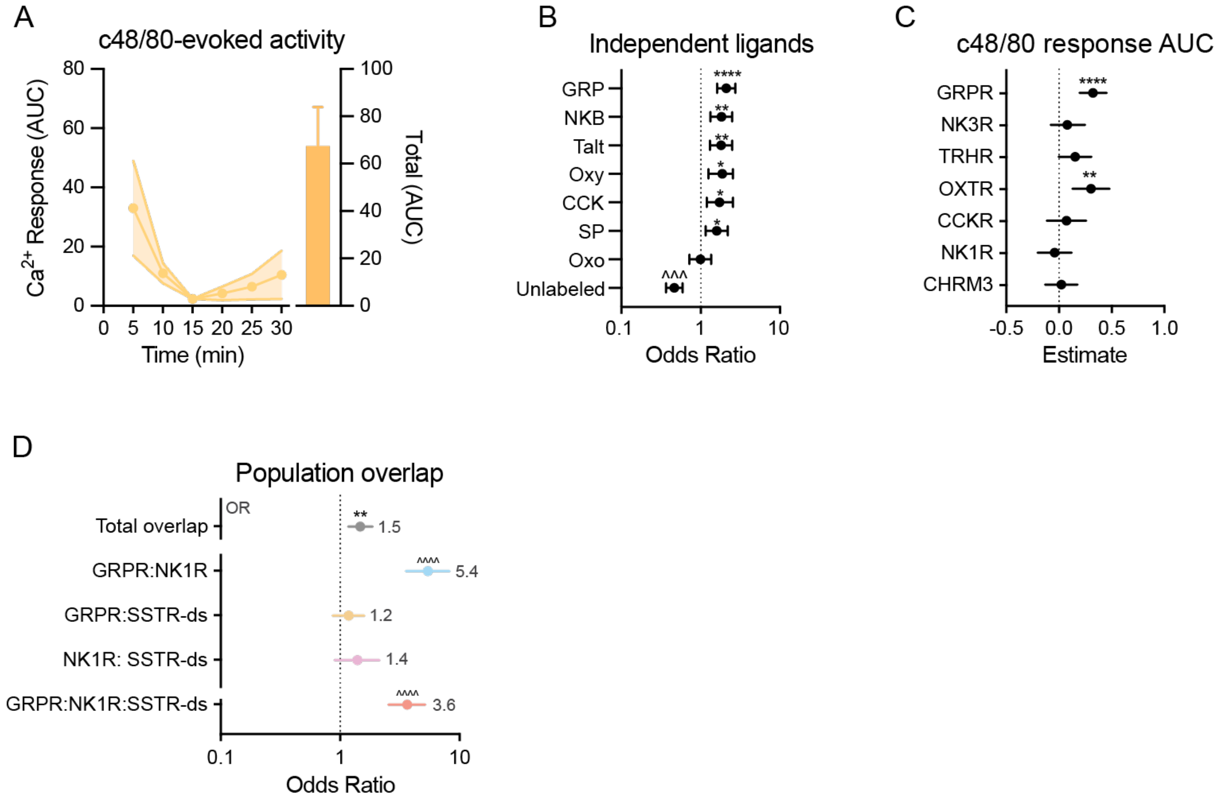


**Supplemental Figure 3**

S3A) AUC over time (left y-axis) and cumulative AUC (right y-axis) following compound 48/80 injection (n=85-325 compound 48/80-responsive neurons/mouse, N=3 mice). Data are shown as mean ± SEM.

S3B) Neurons that respond to GRP, NKB, Talt, Oxy, CCK, and SP are more likely to respond to compound 48/80. (n=85-325 compound 48/80-responsive neurons/mouse, N=3 mice). Odds ratio (OR) analyses, Bonferroni correction for multiple comparisons. Data are shown as OR estimate ± upper and lower 95% CI. * $p < 0.05$, ** $p < 0.01$, *** $p < 0.001$, **** $p < 0.0001$, ^^^ $p < 10^{-8}$.

S3C) GRPR expression is most strongly associated with the magnitude (AUC) of compound 48/80-evoked responses (n=85-325 compound 48/80-responsive neurons/mouse, N=3 mice). Linear regression analysis, Bonferroni correction for multiple comparisons. Data are shown as estimate ± upper and lower 95% CI. ** $p < 0.01$, **** $p < 0.0001$.

S3D) GRPR, NK1R, and SSTR-ds populations overlap at frequencies much higher than expected by the prevalence of each cell type (n=1379 total neurons, 359-610 neurons/mouse, N=3 mice). Odds ratio (OR) analyses, Bonferroni correction for multiple comparisons. OR values for each cell type are in gray. Data are shown as OR estimate ± upper and lower 95% CI. ** $p < 0.01$, ^^^^$p < 10^{-10}$.

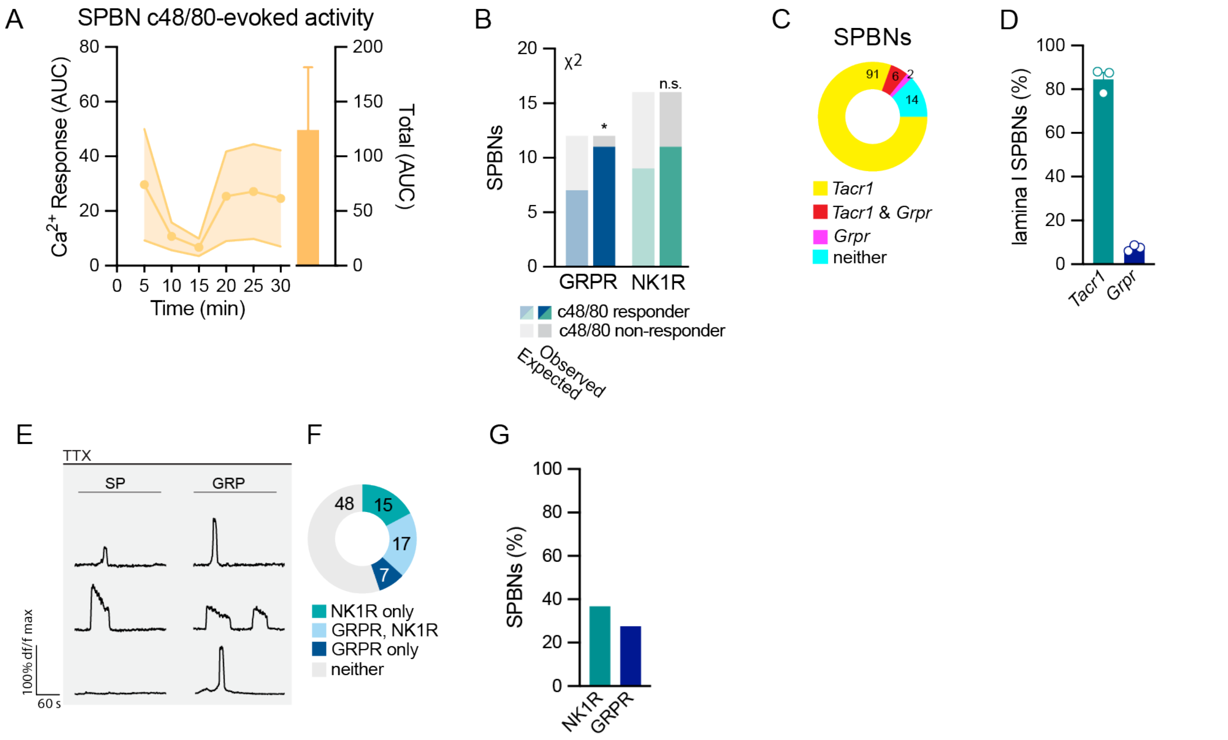


**Supplemental Figure 4**

S4A) AUC overtime (left y-axis) and cumulative AUC (right y-axis) of $Ca^{2+}$ responses in SPBNs following compound 48/80 injection (n=1-10 compound 48/80-responsive SPBNs/mouse, N=3 mice). Data are shown as mean ± SEM.

S4B) Number of GRPR- and NK1R-expressing SPBNs expected to respond to compound 48/80 based on the proportion of all SPBNs that express each receptor (GRPR, 41%; NK1R 55%), versus the observed number of neurons from each population that responded to compound 48/80 (n=29 total SPBNs pooled from N=3 mice). $\chi^2$ test, Bonferroni correction for multiple comparisons. *$p < 0.05$.

S4C) *Grpr* mRNA is expressed in a subset of SPBNs, and the majority of *Grpr* SPBNs also express *Tacr1* (n=114 neurons pooled from N=3 mice).

S4D) Percentage of lamina I SPBNs that express *Tacr1* versus *Grpr* mRNA in C57BL/6 mice. (n=23-50 SPBNs/mouse, N=3 mice). Data are shown as mean ± SEM, with open circles representing individual mice.

S4E) Representative $Ca^{2+}$ imaging traces of SPBNs responding to SP and/or GRP in the presence of TTX.

S4F) GRPR is functionally expressed in a subset of SPBNs, and the majority of GRPR SPBNs also functionally express NK1R (n=87 SPBNs pooled from N=16 mice).

S4G) Percentage of SPBNs that functionally express NK1R versus GRPR in *Vglut2-GCaMP6s* mice (n=87 SPBNs pooled from N=16 mice).

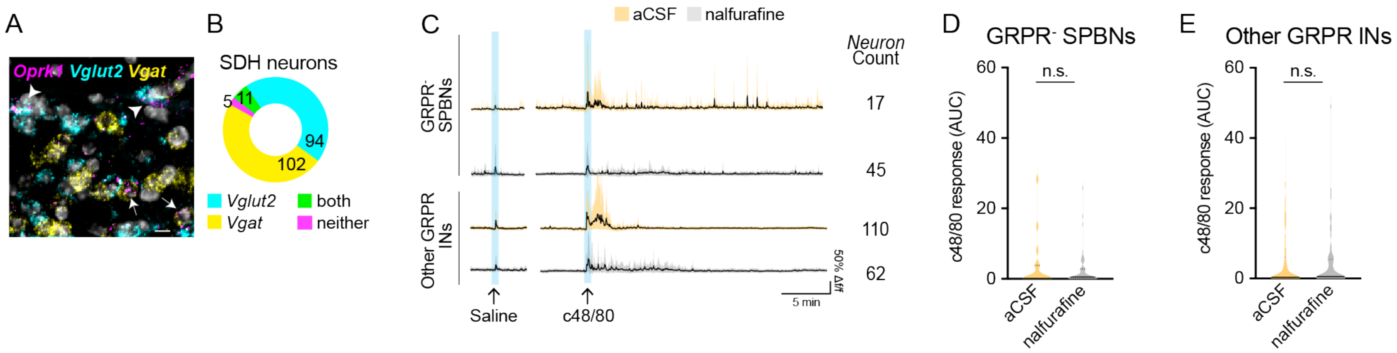


**Supplementary Figure 5**

S5A) Representative image showing expression of *Vglut2* and *Vgat* within *Oprk1* superficial dorsal horn (SDH) neurons. Scale bar, 10 um.

S5B) Quantification of *Vglut* versus *Vgat* expressing *Oprk1* neurons in the SDH (n=212 *Oprk1* neurons pooled from N=4 mice).

S4C) Representative ΔF/F $Ca^{2+}$ traces showing responses of $GRPR^{-}$ SPBNs and other GRPR INs (those that don't respond to additional itch peptides) to intradermal compound 48/80 following pretreatment with aCSF versus nalfurafine, as well as the number of neuron of subtype per condition (neurons pooled from N=3-4 mice per condition). Blue bars correspond to the brief $Ca^{2+}$ transients evoked by intradermal injection alone. Data shown as mean ± 90th percentile.

S4D and S5E) Nalfurafine had no effect on compound 48/80 responses of GRPR INs that do not respond to itch peptides ("Other GRPR INS") or SPBNs that lack GRPR ($GRPR^{-}$ SPBNs: aCSF, n=17 neurons pooled from N=3 mice; nalfurafine, n=45 neurons pooled from N=3 mice. Other GRPR INs: aCSF, n=1178 neurons pooled from N=4 mice; nalfurafine, n=1253 neurons pooled from N=4 mice). Linear mixed-effect model.

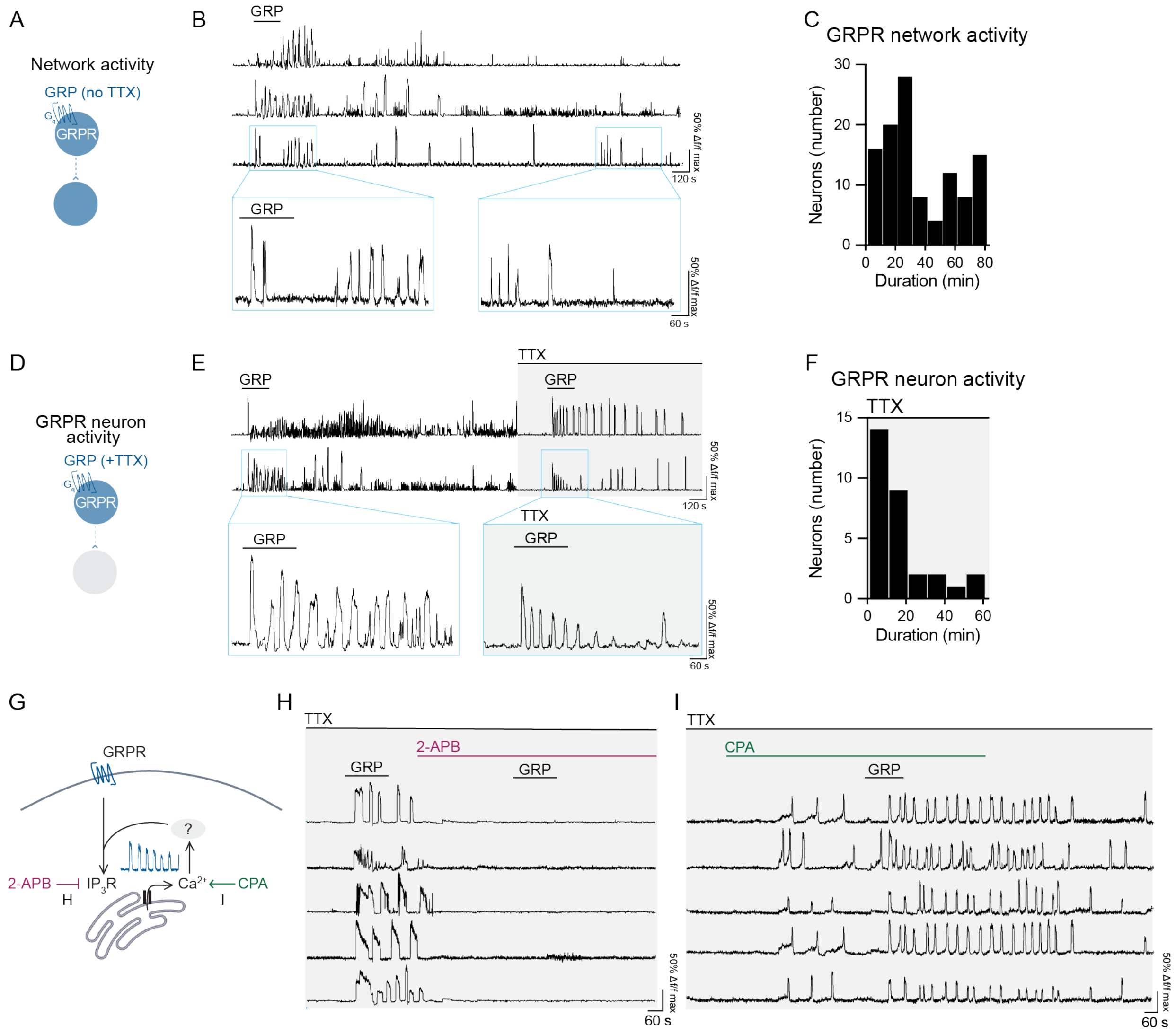


**Supplemental Figure 6**

S6A-B) In one experiment, GRP-evoked activity in the absence of TTX was recorded for 80 min. Representative ΔF/F $Ca^{2+}$ traces from neurons showing GRP-induced repeated $Ca^{2+}$ transients that persist long after GRP is washed out.

S6C) Duration (min) of GRP-evoked network activity (n=111 neurons from N=1 mouse).

S6D-E) In another experiment, GRP-evoked activity in the presence of TTX was recorded for 60 min. Representative ΔF/F $Ca^{2+}$ traces from neurons showing that GRP application evokes prolonged oscillations in both the presence of TTX (500 nM, 8 min pretreatment).

S6F) Duration of GRP-evoked activity in the presence of TTX (n=30 neurons from N=1 mouse).

S6G) Model of $Ca^{2+}$ oscillation in GRPR neurons. Activation of GRPR by GRP opens $IP_3R$, allowing $Ca^{2+}$ influx from internal ER stores. Through unknown feedback mechanisms, this rise in $Ca^{2+}$ sustains the periodic opening of $IP_3R$. 2-APB inhibits the activation of $IP_3R$, and CPA increases intracellular $Ca^{2+}$ level by inhibiting reuptake.

S6H) Representative ΔF/F $Ca^{2+}$ traces from neurons showing $Ca^{2+}$ oscillations in response to GRP application that cease when 2-APB (100 μM) is applied. In the presence of 2-APB, GRP also failed to induce $Ca^{2+}$ responses. Traces representative of N=2 mice.

S6I) Representative ΔF/F $Ca^{2+}$ traces from neurons showing oscillation in response to CPA (10 μM) and increased oscillation frequency when GRP is applied. Traces representative of N=1 mouse.
